# Combined antagonism of Oncostatin M (OSM) and Interleukin 6 (IL-6) provides both anti-fibrotic and anti-inflammatory benefit in pulmonary fibrosis

**DOI:** 10.64898/2025.12.25.695625

**Authors:** Rojo A. Ratsimandresy, Emma Doran, Hart S. Dengler, Sarah E. Headland, Christopher J. Wedeles, Riccardo Guidi, Daqi Xu, Jiatong Liu, Yongchang Shi, Jianyong Wang, Salil Uttarwar, Alexander R. Abbas, Tianhe Sun, Daryle J. DePianto, Katrina B. Morshead, Joe Arron, Jason R. Rock, Jack Bevers, Rajbharan Yadav, Thirumalai Ramalingam, Xia Gao, Hans D. Brightbill, Claire L. Emson, Surinder Jeet, Alexander Arlantico, Aaron Wong, Sherman Yu, Tiffany Hao Tran, Laurie Leong, Monika Dohse, Cary Austin, Patrick Caplazi, Reynold A. Panettieri, Cynthia Koziol-White, William F. Jester, Andrey Shaw, Shimrit Avraham, Hari Menon, Spyros Darmanis, Zora Modrusan, Kimberly Kajihara, Masakazu Kanamori, Hideaki Shimada, Nathaniel R. West, Gerald Nakamura, Jason A. Vander Heiden, Mark S. Wilson

## Abstract

Interstitial lung diseases (ILDs), including idiopathic pulmonary fibrosis (IPF) and systemic sclerosis-associated ILD (SSc-ILD), are irreversible fibrosing diseases with a mean survival time of less than 5 years for IPF. Poorly understood etiology and complex pathogenesis of these diseases have hampered the identification and development of effective therapeutics. Existing treatments can slow progressive fibrosis and lung function decline but do not stop it entirely, resulting in a minimal impact on patient survival. Thus, novel therapeutic interventions are needed. Tocilizumab, an anti–IL-6 receptor antibody, was recently approved by FDA for the treatment of SSc-ILD based on evidence demonstrating a reduction in the rate of lung function decline. In this study, we have characterized an IL-6-driven feed-forward myeloid axis contributing to lung inflammation providing a mechanistic hypothesis for tocilizumab. Concomitantly we have identified an oncostatin M (OSM) orchestrated lung injury response contributing to epithelial and endothelial cell disruption, myofibroblast activation and fibrosis. Despite dysregulated expression of *IL11* in IPF, SSc-ILD, and murine models of fibrosis, we found no evidence for a pro-fibrotic role for IL-11 *in vitro* or *in vivo*. Instead, in pre-clinical models of IPF we demonstrate that antagonism of OSM alone, or to a greater degree in combination with IL-6, reduced lung fibrosis and inflammation. Translating these observations, we validated gp130:OSMR, rather than gp130:LIFR, as the dominant human receptor complex used by OSM, identifying OSMR as a potential therapeutic target to stall fibrosis.

## Introduction

Interstitial lung diseases (ILDs), including idiopathic pulmonary fibrosis (IPF) and systemic sclerosis-associated ILD (SSc-ILD), are chronic, irreversible and often fatal pulmonary diseases that can carry an average prognosis of less than 5yrs^1–3^. IPF and up to 80% of SSc patients^4^ present with pulmonary inflammation, vascular dysfunction, accumulation of subpleural fibroblasts and progressive fibrosis. Despite a good understanding of the pathophysiology of ILDs, there are very few treatment options available. Many prognostic biomarkers have been identified, including interleukin 6 (IL-6). Serum levels of IL-6 are elevated in patients with IPF and SSc and are predictive of lung function decline^5^. Experimental data support a pathogenic role for IL-6 with deletion of *Il6* or blocking the IL-6 pathway with monoclonal antibodies (mAbs) protecting mice from bleomycin (BLM)-induced lung and skin fibrosis, although the mechanism of protection has not been fully elucidated^6–8^. These observations spurred the clinical investigation of IL-6 in pulmonary fibrosis using tocilizumab (TCZ), a humanized monoclonal antibody directed against IL-6R that inhibits IL-6 signaling and previously approved for the treatment of rheumatoid arthritis, juvenile idiopathic arthritis, and giant cell arteritis^9^. Phase II (FaSScinate) and III (FocuSSced) clinical trials in patients with SSc found significantly fewer patients in the TCZ arm experienced a decline in lung function over 48 weeks of treatment, compared to placebo^10,11^. In 2019, TCZ was approved by the FDA for the treatment of SSc-ILD; however, these studies did not meet their primary endpoints of reducing skin fibrosis^10,11^. To date, pirfenidone, nintedanib, and nerondomilast are the only approved antifibrotic therapies for ILD’s to slow pulmonary function decline^12,13^, but they have well-characterized tolerability concerns and only moderately improve patient survival^14,15^,thus leaving a need for more effective therapeutic options to prevent progressive fibrosis and improve patient outcomes.

Several profibrotic mediators have been described, including growth factors (CTGF, TGFβ family members), cytokines (IL-13 and IL-11), integrins, and vasoactive peptides^16^. In addition to these, oncostatin M (OSM) has long been associated with fibrosis^17–19^ and is elevated in the bronchoalveolar lavage (BAL) fluid of SSc-ILD and IPF patients^20^. In pre-clinical studies, elevated OSM has accompanied fibrosis in a variety of organs^21,22^ and can induce fibrotic responses when exogenously delivered or over-expressed in mice^20,23–28^. OSM-driven fibrosis appears to be independent of RAG-dependent T and B cells, cKIT-expressing mast cells and basophils, STAT6^26^ and IL-4R-dependent IL-4 and IL-13 or TGFβ family members^20^, suggesting a non-overlapping and independent OSM-mediated mechanism of fibrosis. Despite these studies indicating that OSM is sufficient to drive fibrosis, the role of OSM in pulmonary fibrosis and the benefit of antagonizing OSM in pre-clinical or clinical settings is currently unclear.

Here we describe a mechanism of how IL-6 contributes to pulmonary fibrosis, identify a critical role for OSM contributing to fibrosis, and demonstrate how antagonism of OSM alone, or combined with IL-6, can provide therapeutic benefit. Specifically, we show that IL-6 promotes the recruitment of a CD64^+^ monocyte-derived macrophage population, providing a mechanistic explanation for the benefit observed with IL-6R antagonism in the clinic.

IL-6 signals through a heterodimeric receptor complex composed of IL6Rα and gp130. Several other cytokines that have been implicated as mediators of pathological fibrosis, including IL-11 and OSM, also use gp130 as a coreceptor. With the aim of identifying pro-fibrotic mediators that can be targeted to stall progressive fibrosis we initially assessed the contribution of IL-11 and OSM. We did not find any evidence that IL-11 contributes to pulmonary, kidney or liver fibrosis and instead identified a non-redundant and critical role for OSM. Furthermore, IL-6 and OSM contribute in a non-overlapping manner, with combined antagonism of IL-6 and OSM providing superior protection from lung injury, inflammation and fibrosis. Translating these observations to human systems, we identify *OSMR* expressing fibroblasts, epithelial, and endothelial cells in both murine and human IPF lung tissue, characterize OSM-driven signaling, transcriptional and physiological responses in all three primary human cell types, and support OSMR as a viable therapeutic target to prevent progressive fibrosis.

## Results

### IL-6 drives CD64^+^ macrophage activation in murine models of pulmonary inflammation and fibrosis

We have clinically validated an important role for IL-6 signaling in lung function decline in patients with SSc-ILD^11^, however, IL-6R antagonism only partially impacted fibrosis endpoints and fibrosis-related biomarkers. To better understand the mechanisms of IL-6, we first investigated the role of IL-6 in the murine bleomycin (BLM)-induced lung injury, inflammation, and fibrosis model. Confirming previous reports of antagonizing this pathway ^6–8^, *Il6r^−/−^* mice had reduced lung injury and inflammation at both day 8 and day 24 post-BLM (**Fig. 1a**) with reduced total hydroxyproline, an amino acid necessary for collagen biosynthesis (**Fig. 1b**), along with reduced *Col1a1* and *Col1a2* gene expression in lung tissue (**Fig. 1c**). Correlating with reduced disease in *Il6r^−/−^* mice was a reduction in the proportion and total number of CD64^+^SiglecF^−^ macrophages (CD45^+^CD11c^+^SiglecF^−^MHCII^+^CD11b^+^CD64^+^) at both day 8 and day 24 post-BLM (**Fig. 1d**), suggesting that IL-6 contributes to macrophage recruitment to the fibrotic lung. To better characterize lung macrophages at these early time points, we isolated CD64^+^ (CD45^+^CD11c^+^ Siglec F^−^MHCII^+^CD11b^+^CD64^+^) and alveolar (CD45^+^CD11c^+^Siglec F^+^) macrophages from the lungs of BLM-treated WT mice at day 5 and 14 for both transcriptional and secretome analysis. At day 5 post-BLM and to lesser extent at day 14, CD64^+^ macrophages secreted large amounts of inflammatory monocyte-recruiting chemokines, CCL2, CCL3, and CCL4 (**Extended data 1a**), suggesting that IL-6-dependent CD64^+^ macrophages could contribute, in a feed forward manner, to monocyte trafficking into the site of injury. Supporting this notion, deeper transcriptional analysis of *ex vivo* CD64^+^, but not alveolar, macrophages identified elevated transcripts encoding a variety of inflammatory monocyte markers, cytokines, and chemokines, including *Ccl8*, *Ccl12*, *Cx3cr1*, *Mmp13,* and *complement*. Of note, these characteristic features have all been identified in a variety of previously reported fibrogenic macrophages^29–31^ (**Fig. 1e**). To determine whether IL-6 regulated the activation state and recruitment of monocytes into the lung, we primed murine bone marrow-derived macrophages with IL-4/IL-13 or IFNγ/LPS before exposing them to a pulse of IL-6. As expected, IL-4/IL-13 up-regulated *Arg1* and *Cd206*, and, with the addition of IL-6, these levels doubled, unlike IFNγ/LPS primed macrophages where *Nos2* expression was unchanged with the addition of IL-6 (**Extended data 1b, 1c**). Most striking was the observation that IL-6 dramatically increased both transcription and secretion of CCL2 from IL-4/IL-13 primed macrophages (**Extended data 1d**), suggesting that IL-6 may be a potent activating signal for fibrogenic macrophages in the lung, driving the secretion of CCL2 and subsequent recruitment of monocytes.

**Figure 1.**
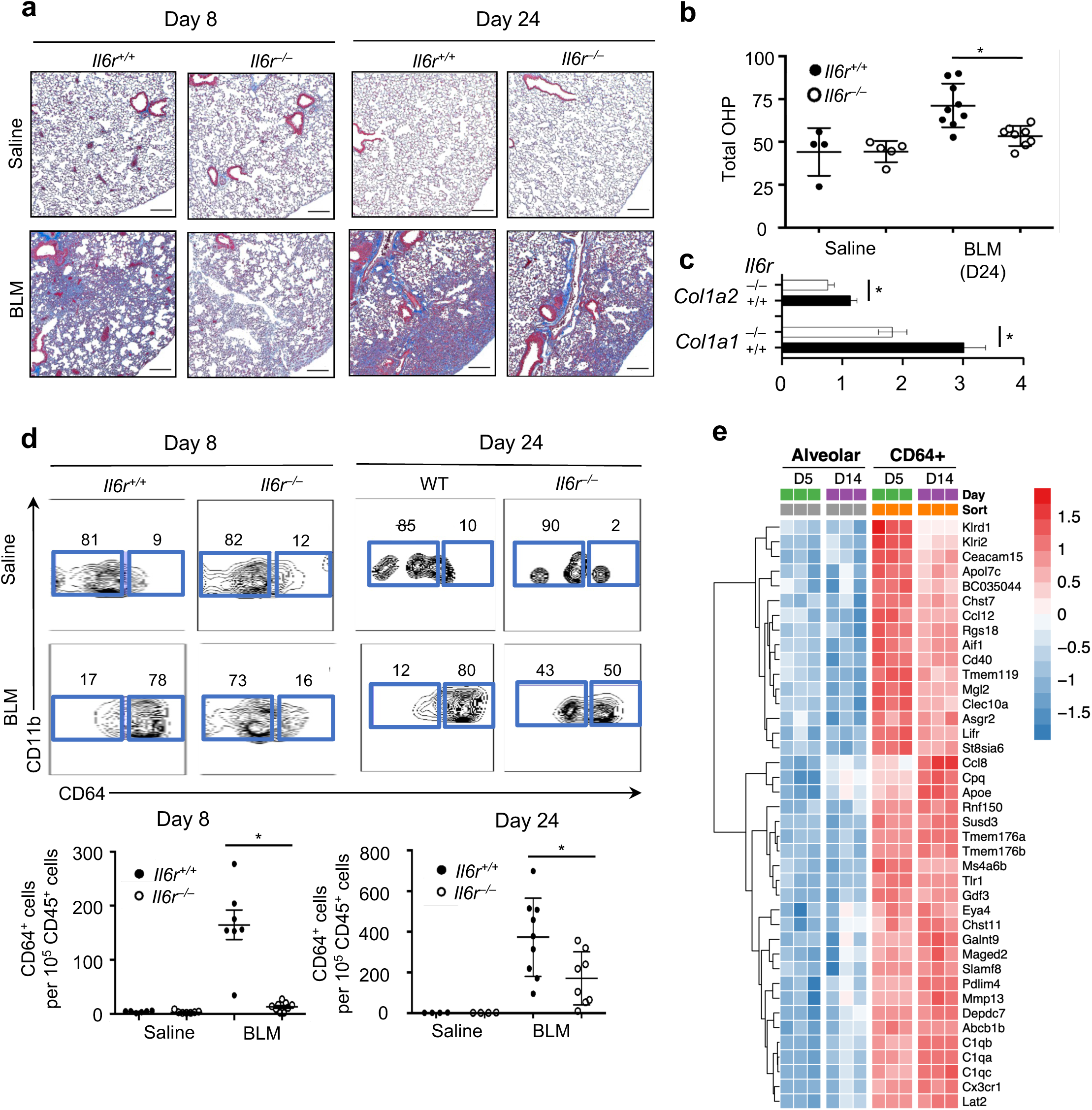
IL-6 dependent CD64^+^ macrophage populations drive fibrotic disease. a. WT (*Il6r*^+/+^) and IL-6-deficient (*Il6r*^−/−^) mice were given intratracheal saline or bleomycin (BLM) on day 0, 2 and 4, as described in methods. Masson’s Trichrome stained lung tissue at day 8 and day 24 post BLM was carried out to identify collagen deposition. 4-8 mice per group, representative images shown. b. Total hydroxyproline (Total OHP, ug/half lung) lung tissue was measured at day 24. 4-8 mice per group. P value calculated by t-test. Graphs show mean ± SD. * P<0.05. c. Gene expression measured in lung tissue by qRT-PCR. 4-8 mice per group. P value calculated by t-test. Graphs show mean ± SD. * P<0.05. d. FACS analysis of lung tissue at day 8 and day 24 from saline or BLM treated WT (*Il6r*^+/+^) and IL-6-deficient (*Il6r*^−/−^) mice showing representative FACS plots (top) and numbers of CD64^+^ macrophages (CD45^+^CD11c^+^SigF^+^MHCII^+^CD11b^+^CD64^+^) per 10^5^ CD45^+^ cells (bottom). 4-8 mice per group. P value calculated by t-test. Graphs show mean ± SD. * P<0.05. e. Heatmap showing the top 40 (by fold-change) differentially expressed genes (DEG) in alveolar macrophages (CD45^+^CD11c^+^SigF^−^) and CD64^+^ macrophages (CD45^+^CD11c^+^SigF^−^MHCII^+^CD11b^+^CD64^+^) macrophages from WT mice treated with saline or BLM at day 5 or day 14.

Transcriptional analysis of IL-6-treated macrophages identified a suite of transcripts associated with inflammatory monocytes^32^ and pro-fibrotic mediators, including *Il4ra, Vegfd, Serpina3g, Ccl2, Ccl7,* and *Ccl24* (**Extended data 1e**). To determine whether these observations in murine macrophages translated to human macrophages, we primed human monocyte-derived macrophages with IL-4/IL-13 and treated them with IL-6. As expected, IL-6 significantly increased CCL18 transcripts and secretions, a chemokine known to be prognostic for worse ILD^33,34^ (**Extended data 1f**). Broader analysis of the differentially expressed genes revealed that IL-6 also amplified expression of several additional fibrosis-associated genes including *ENPP2*, *DHRS2*, *COL24A1*, *CXCL1*, *FZD2*, and *IL17RB* (**Extended Fig 1g**). Following these mechanistic and *in vitro* observations in macrophages, we reanalyzed IPF and SSc-ILD patient samples and observed that IL-6-regulated macrophage genes *FCGR1A (CD64)*, *CCL2,* and *CCL18* were all increased in the lung tissue of IPF (**Extended data. 2a**) or skin of SSc-ILD (**Extended data. 2b**) patients. Following 24 weeks of TCZ treatment, there was a very clear and strong pharmacodynamic effect on serum levels of CCL18 in the faSScinate trial (**Extended data. 2c**), suggesting that IL-6-regulated macrophage activation in these patients may be reduced. Thus, IL-6 may contribute to lung function decline in patients with ILD via an inflammatory macrophage-mediated activation pathway.

### Redundant, pathogenic, and protective roles of IL-11 in the lung, liver and kidney

A closely related gp130-dependent cytokine, IL-11, has recently emerged as an important contributor to lung, liver, and cardiac fibrosis^21,35–38^. We therefore profiled expression of *IL11* in a variety of tissues and found it to be significantly increased in the lung tissue of IPF patients (**Fig. 2a**), though not the skin of SSc patients (**Fig. 2b**).

**Figure 2.**
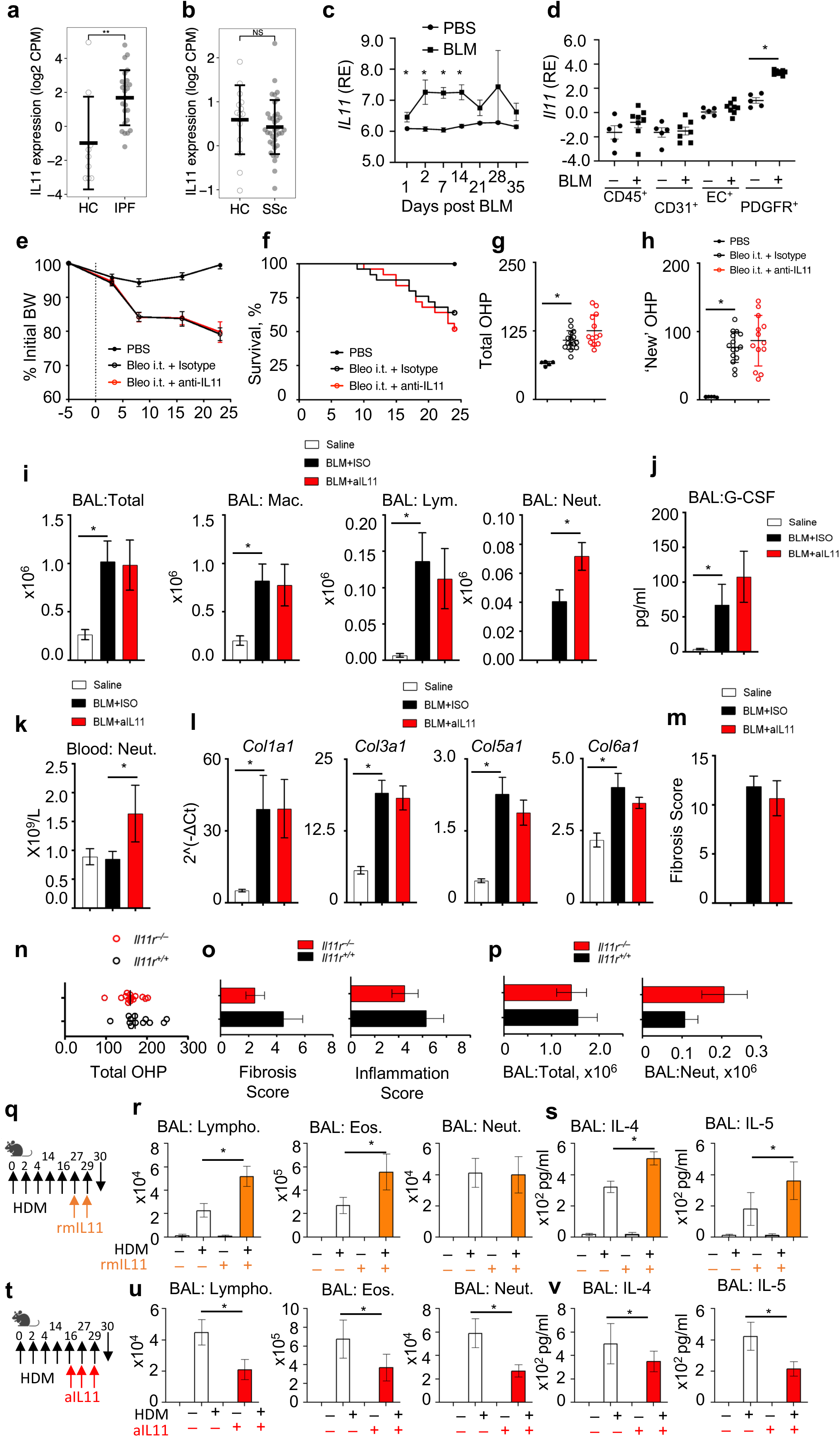
Redundant, pathogenic and protective roles of IL-11 in lung, liver and kidney. a. Lung biopsies were obtained from healthy controls (n=9) and IPF patients (n=24), RNA-seq was carried out and *IL11* transcripts analyzed. Graphs show log_2_ normalized CPM, mean ± SD. * FDR P<0.05, ** FDR P<0.005; NS not significant. b. Skin biopsies obtained from healthy controls (n=14) and SSc patients (n=36), participating in the focuSSced clinical trial, RNA-seq was carried out and *IL11* transcripts analyzed. Graphs show log_2_ normalized CPM, mean ± SD. * FDR P<0.05, ** FDR P<0.005; NS not significant. c. WT C57BL/6J mice were given intratracheal saline (PBS) or bleomycin (BLM) on day 0, 2 and 4, as described in methods. Lung tissue was collected on day 1, 2, 7, 14, 21, 28 and 35 post BLM. RNA was extracted and *Il11* transcripts determined by RNA sequencing. N = 5-25 mice per group. Graphs show mean ± SD. * P<0.05 at each time point. d. WT C57BL/6J mice were given intratracheal saline (PBS) or bleomycin (BLM) on day 0, 2 and 4, as described in methods. Lung tissue was collected on day 24 post BLM. Lung tissue was dissociated for FACS staining and sorting. CD45^+^ CD31^+^, Epcam^+^ and PDGFR^+^ cells were purified, with RNA extracted for RNA sequencing. N = 3, 5-10 mice per pooled group. Graphs show mean ± SD. * P<0.05. e. WT C57BL/6J mice were given intratracheal saline (PBS) or bleomycin (BLM) on day 0, 2 and 4, as described in methods. Mice were given anti-IL-11 mAb (MAB418, 20mg/kg every 3 days, from day −1). Body weight was monitored weekly. f. Survival was monitored and mice with >25% weight loss were euthanized. g. Total hydroxyproline (Total OHP, ug/half lung) in lung tissue was measured at day 24. 5-25 mice per group. P value calculated by t-test. Graphs show mean ± SD. * P<0.05. h. ‘New’ hydroxyproline (New OHP, ug/half lung) in lung tissue was measured in deuterated water treated mice, as described in methods, at day 24. 5-25 mice per group. P value calculated by t-test. Graphs show mean ± SD. * P<0.05. i. Bronchoalveolar lavage (BAL) was performed on mice at day 24, as described in methods. Total and differential cell counts (Macrophages, Mac; Lymphocytes, Lym; Neutrophils, Neut) were determined from each mouse. 5-25 mice per group. P value calculated by t-test. Graphs show mean ± SD. * P<0.05. j. Bronchoalveolar lavage (BAL) was performed on mice at day 24, as described in methods. GCSF was measured by Luminex. 5-25 mice per group. P value calculated by t-test. Graphs show mean ± SD. * P<0.05. k. Whole blood was collected from mice at day 24. WBC and differential cell analysis was measured on a Cobas system. 5-25 mice per group. P value calculated by t-test. Graphs show mean ± SD. * P<0.05. l. Gene expression measured in lung tissue by qRT-PCR. 5-25 mice per group. P value calculated by t-test. Graphs show mean ± SD. * P<0.05. m. Lung pathology (fibrosis score) was assessed in a blinded manner, as described in methods. n. WT (*Il11r*^+/+^) and IL-11R-deficient (*Il11r*^−/−^) C57BL/6J mice were given intratracheal saline (PBS) or bleomycin (BLM) on day 0, 2 and 4, as described in methods. Total hydroxyproline (Total OHP, ug/half lung) in lung tissue was measured at day 24. 10 mice per group. P value calculated by t-test. Graphs show mean ± SD. * P<0.05. o. Lung pathology (fibrosis score and inflammation score) was assessed in a blinded manner, as described in methods. 10 mice per group. P value calculated by t-test. Graphs show mean ± SD. * P<0.05. p. Bronchoalveolar lavage (BAL) was performed on mice at day 24, as described in methods. Total and neutrophil cell counts were determined from each mouse. 10 mice per group. P value calculated by t-test. Graphs show mean ± SD. * P<0.05. q. C57BL/6 mice were treated with 10ug HDM in 25ul via the intratracheal route on day 0, 2, 4, 14, 16, 27, 29, 30. Mice were also given rmIL-11 via the intratracheal route, as indicated and described in methods, on day 27 and 29. Mice were analyzed for airway inflammation on day 30. r. Bronchoalveolar lavage (BAL) was performed on mice at day 24, as described in methods. Total and differential cell counts (Lymphocytes, Lym; Neutrophils, Neut; Eosinophils, Eos) were determined from each mouse. 5-15 mice per group. P value calculated by t-test. Graphs show mean ± SD. * P<0.05. s. Bronchoalveolar lavage (BAL) was performed on mice at day 30, as described in methods. IL-4 and IL-5 was measured by Luminex. 5-15 mice per group. P value calculated by t-test. Graphs show mean ± SD. * P<0.05. t. C57BL/6 mice were treated with 10ug HDM in 25ul via the intratracheal route on day 0, 2, 4, 14, 16, 27, 29, 30. Mice were also given anti-mouse IL-11 mAbs i.p., as indicated and described in methods, on day 16, 27 and 29. Mice were analyzed for airway inflammation on day 30. u. Bronchoalveolar lavage (BAL) was performed on mice at day 24, as described in methods. Total and differential cell counts (Lymphocytes, Lym; Neutrophils, Neut; Eosinophils, Eos) were determined from each mouse. 5-15 mice per group. P value calculated by t-test. Graphs show mean ± SD. * P<0.05. v. Bronchoalveolar lavage (BAL) was performed on mice at day 30, as described in methods. IL-4 and IL-5 was measured by Luminex. 5-15 mice per group. P value calculated by t-test. Graphs show mean ± SD. * P<0.05.

Similarly, in the lung tissue of mice treated with BLM, *Il11* was elevated from as early as day 2 (**Fig. 2c**). More specifically, purifying a variety of cells from the lung tissue of mice, *Il11* was predominantly expressed in PDGFR^+^ fibroblasts at steady state and significantly increased following BLM treatment (**Fig. 2d**). Encouraged by the *Il11* expression profile in both human and murine fibrotic lung tissue, we either generated or sourced recombinant murine and human IL-11 and anti-IL-11 blocking mAbs to test the function and role of IL-11. We first validated that IL-11 (rmIL11, rhIL11) could activate primary and fibroblast cell lines (**Extended data 3a,** ED_50_ = 15-70 ng/ml) and that anti-IL-11 blocking mAbs could both bind to IL-11 (**Extended data 3b**) and block rmIL-11-mediated fibroblast activation *in vitro* (**Extended data 3c**, IC_50_ = 33.4 nM).

Finally, we ensured that anti-IL-11 mAbs could inhibit IL-11-driven pSTAT3 *in vivo* and had a favorable pharmacokinetics profile (T_½_ = 15.8 days) (**Extended data 3d, 3e**). To test whether IL-11 contributed to BLM-induced lung inflammation and fibrosis, we dosed mice with 20 mg/kg of anti-IL-11 every week, prior to and following BLM treatment to ensure complete pathway blockade. BLM treated mice lost weight and succumbed to BLM exposure irrespective of IL-11 antagonism (**Fig. 2e, f**). Furthermore, elevated levels of total (**Fig. 2g**) or newly synthesized (**Fig. 2h**) hydroxyproline (from day 9 – day 24), was unchanged following IL-11 blockade. Similarly, increased airway infiltrates and GCSF following BLM treatment were unchanged following IL-11 blockade (**Fig. 2i, j**).

Interestingly, we observed a small increase in circulating neutrophils following IL-11 inhibition (**Fig. 2k**). Finally, gene expression in lung tissue (**Fig. 2l**) and lung pathology (**Fig. 2m**) confirmed that in our hands IL-11 appears to be redundant for BLM-induced lung damage, inflammation, and fibrosis. To test the IL-11 pathway in a different way, we sourced *Il11r*^−/−^ mice and confirmed that lung fibroblasts from these mice were unresponsive to rmIL-11, but responsive to rmIL-11 fused to rmIL-11 receptor (IL-11:IL-11R) (**Extended data 3f, 3g**). Similar to the pharmacological approach with anti-IL-11 mAbs, *Il11r*^−/−^ mice developed pulmonary inflammation and fibrosis similar to WT (*Il11r*^+/+^) mice, with comparable lung hydroxyproline content (**Fig. 2n**), fibrosis, and tissue inflammation (**Fig. 2o**), and with a similar degree of airway inflammation (**Fig. 2p**). To more directly test whether rmIL-11 could activate lung fibroblasts, we cultured WT murine lung-derived fibroblasts with rmIL-11 and assessed both inflammatory cytokine secretion and ACTA2 protein expression. Although rmIL-11 could induce a variety of cytokines, rmIL-11 failed to induce ACTA2 expression (**Extended data 3h, 3i**). Previous studies have also implicated IL-11 in the pathogenesis of obstructive airway disease, such as asthma, in both human and murine studies^39–42^. We therefore took a gain- and loss-of-function approach and confirmed that rmIL-11 given to house dust mite (HDM) exposed mice could enhance lymphocytic and eosinophilic airway inflammation (**Fig. 2q, 2r**), with a concordant increase in BAL IL-4 and IL-5 (**Fig. 2s**). In agreement with an important role for IL-11 contributing to airway inflammation, anti-IL-11 treated mice had significantly reduced cellular airway infiltration and BAL cytokine secretions (**Fig. 2t, 2u, 2v**). Collectively, these studies indicate that while IL-11 contributes to airway inflammation *in vivo* and that IL-11 can induce inflammatory cytokine responses in lung fibroblasts *in vitro*, IL-11 does not appear to have fibrogenic properties in the lung *in vivo* or with lung-derived fibroblasts *in vitro*. To further explore the role of IL-11 in other fibrotic responses, we extended our studies into the kidney using two different kidney fibrosis models, a nephrotoxic serum (NTS)-induced kidney damage and fibrosis model and a unilateral ureteral obstruction (UUO) model of kidney fibrosis. NTS treated *Il11r*^+/+^ and *Il11r*^−/−^ mice both developed similar degrees of glomerular damage and fibrosis (**Extended data 4a, 4b)**. Similarly, nephropathy in WT mice caused by UUO was similar between mice given control mAbs and anti-IL-11 mAbs, with similar levels of hydroxyproline and expression of *Col1a1*, *Tnfa,* and *Il6* in the kidney (**Extended data 3c, 3d**). Finally, given the reported roles of IL-11 in the liver, we tested whether IL-11 blockade could prevent the choline-deficient, L-amino acid-defined, high-fat diet (CDA-HFD)-induced murine model of nonalcoholic steatohepatitis. Again, anti-IL-11 mAb-treated mice had a similar increase in liver weight and liver damage, with increased AST and ALT (**Extended data 4e**), as mice given control mAbs. Moreover, mice fed a CDA-HFD and given rmIL-11 daily from day 7-11 had similar liver weight increase but significantly less liver damage with reduced serum AST and ALT levels (**Extended data 4f**), indicating that IL-11 has hepato-protective properties as previously indicated^43^. Thus, IL-11 contributes to airway inflammation and protection from CDA-HFD-induced liver injury, but does not appear to contribute to pulmonary or kidney fibrosis. This was also the conclusion from a recent study by Tan and colleagues^44^, who also suggested that IL-11 may not act downstream of TGFβ and that IL-11 may not contribute to fibrogenesis.

### Significant and non-redundant role of OSM in pulmonary fibrosis

OSM has both positively and negatively been associated with fibrosis^19–22,25,27,28,45–47^, but to the best of our knowledge, the requirement for OSM and OSMR signaling in pulmonary fibrosis has not been tested, genetically or pharmacologically. *OSM* was elevated (though not statistically significant) in the lung tissue from IPF patients (**Fig. 3a**) and the skin of SSc patients (**Fig. 3b**). Similarly, *Osm* was significantly elevated in the lung tissue of BLM-treated mice from day 2 onwards (**Fig. 3c**), similar to *Il11* (**Fig. 2d**). However, unlike *Il11*, *Osm* was largely restricted to CD45^+^ cells (**Fig. 3d**). Within CD45^+^ cells, *Osm* was present in both alveolar and CD64^+^ populations and was significantly increased in CD64^+^ macrophages following BLM treatment at day 5 and day 14 (**Fig. 3e**). To formally test the requirement for OSM in BLM-induced pulmonary fibrosis we treated *Osm*^+/+^ and *Osm*^−/−^ mice with BLM. Both groups had similar weight loss (**Fig. 3f**) and survival (**Fig. 3g**) post-BLM; however, *Osm*^−/−^ mice had significantly less lung damage (determined by changes in tissue volume [TV]) (**Fig. 3h**), less total and new hydroxyproline deposited in the lung (**Fig. 3i, 3j**), and significantly less tissue pathology (**Fig. 3k, Extended data 5a**). Interestingly, *Osm*^−/−^ mice had a similar degree of tissue inflammation as *Osm*^+/+^ mice (**Fig. 3l**), suggesting that, in this BLM model, OSM is predominantly pro-fibrotic rather than pro-inflammatory. Encouraged by these genetic studies, we tested whether a pharmacological approach, using an anti-OSM blocking mAb, as previously characterized^48^, would afford similar benefit to WT mice exposed to BLM. Similar to *Osm*^−/−^ mice, BLM-treated WT mice given an anti-OSM blocking mAb had similar weight loss and survival as WT mice given a control mAb (**Fig. 3m, 3n**), but had significantly less total and new hydroxyproline deposited in the lung compared to control mAb treated mice (**Fig. 3o, 3p**). Transcriptional analysis of lung tissue from anti-OSM treated mice identified a variety of fibrotic pathways that were reduced following OSM blockade, including significantly reduced ECM regulators (*Timp1, Mmp10, −12, −13, −14, and −19*) (**Fig. 3q**), collagen synthesizing and regulating genes (*Col1a1, Col1a2, Col3a1, Ereg,* and *Has2*), and, most notably, *Tnc*, which encodes the potent pro-fibrotic hexameric ECM glycoprotein, Tenascin-C^49^ (**Fig. 3r**). Other well-described pro-fibrotic mediators, *Tgfb1, Tgfb2, Tgfb3, Il13,* and *Ctgf* experienced little change following OSM blockade (**Extended data 5b**), suggesting that OSM-driven fibrosis may be independent of these pro-fibrotic pathways. Finally, pathway analysis revealed the extent of the benefit of OSM blockade, with several fibrotic pathways (“wound healing”, “IPF signaling”, “pulmonary healing”, “hepatic fibrosis”) reduced (**Fig. 3s**).

**Figure 3.**
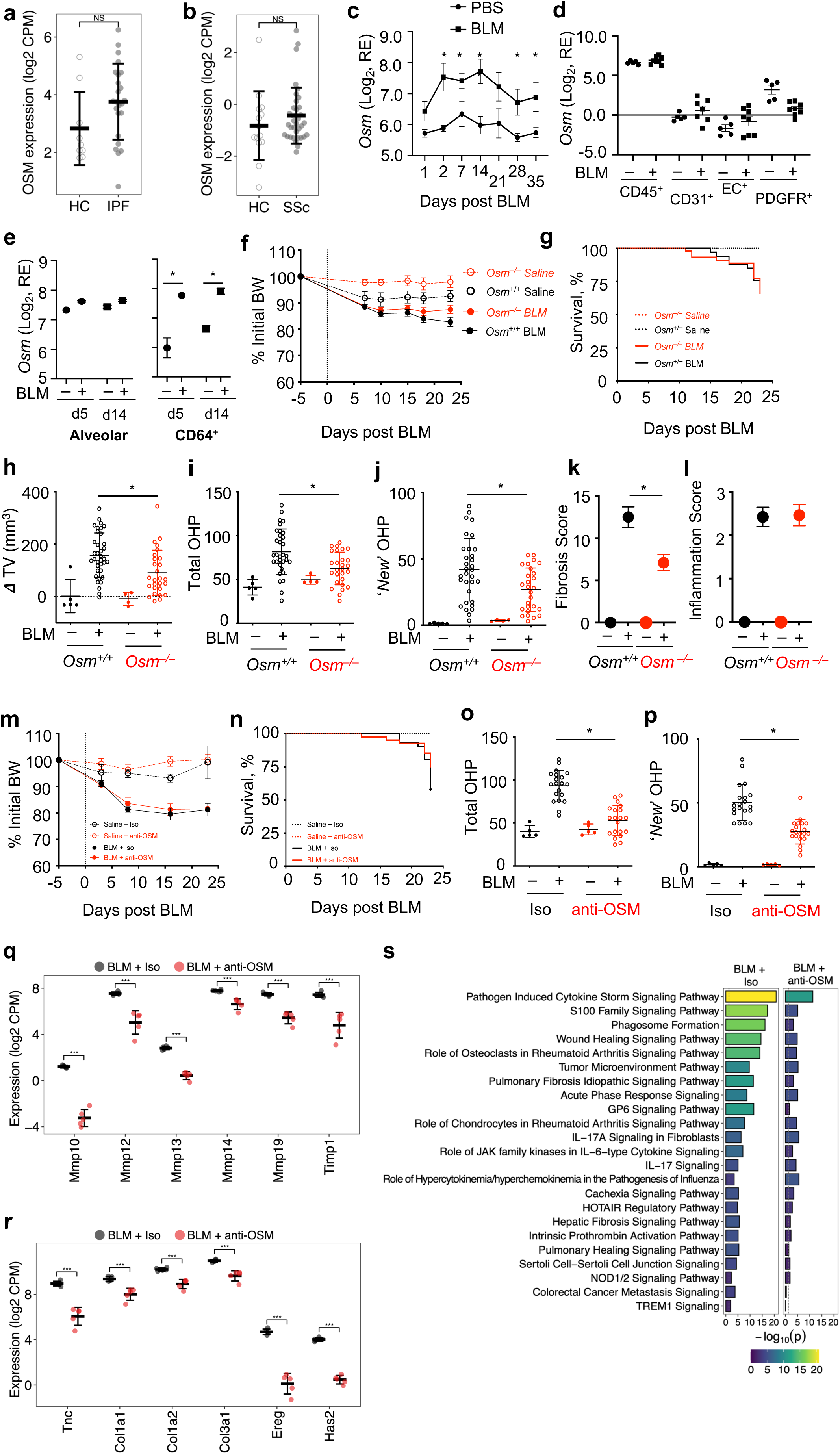
Significant and non-redundant role of Oncostatin M (OSM) in pulmonary fibrosis. a. Lung biopsies were obtained from healthy controls (n=9) and IPF patients (n=24), RNA-seq was carried out and *OSM* transcripts analyzed. Graphs show log_2_ normalized CPM, mean ± SD. * FDR P<0.05, ** FDR P<0.005; NS not significant. b. Skin biopsies obtained from healthy controls (n=14) and SSc patients (n=36) participating in the focuSSced clinical trial and *OSM* transcripts measured. Graphs show log_2_ normalized CPM, mean ± SD. * FDR P<0.05, ** FDR P<0.005; NS not significant. c. WT C57BL/6J mice were given intratracheal saline (PBS) or bleomycin (BLM) on day 0, 2 and 4, as described in methods. Lung tissue was collected on day 1, 2, 7, 14, 21, 28 and 35 post BLM. RNA was extracted and *Osm* transcripts determined by RNA sequencing. N = 5-25 mice per group. Graphs show mean ± SD. * P<0.05 at each time point. d. WT C57BL/6J mice were given intratracheal saline (PBS) or bleomycin (BLM) on day 0, 2 and 4, as described in methods. Lung tissue was collected on day 24 post BLM. Lung tissue was dissociated for FACS staining and sorting. CD45^+^ CD31^+^, Epcam^+^ and PDGFR^+^ cells were purified, with RNA extracted for RNA sequencing. N = 3, 5-10 mice per pooled group. Graphs show mean ± SD. * P<0.05. e. *Osm* expression in alveolar macrophages (CD45^+^CD11c^+^SigF^−^) and CD64^+^ macrophages (CD45^+^CD11c^+^SigF^−^MHCII^+^CD11b^+^CD64^+^) macrophages from WT mice treated with saline or BLM at day 5 or day 14. N = 3, 5-10 mice per pooled group. Graphs show mean ± SD. * P<0.05. f. WT (*Osm*^+/+^) and OSM-deficient (*Osm*^−/−^) C57BL/6J mice were given intratracheal saline (PBS) or bleomycin (BLM) on day 0, 2 and 4, as described in methods. Body weight was monitored weekly and expressed as % of initial weight. g. Survival was monitored and mice with >25% weight loss were euthanized. h. Tissue volume (TV) was determined at day 22, as described in methods. i. Total hydroxyproline (Total OHP, ug/half lung) in lung tissue was measured at day 24. 5-25 mice per group. P value calculated by t-test. Graphs show mean ± SD. * P<0.05. j. ‘New’ hydroxyproline (New OHP, ug/half lung) in lung tissue was measured in deuterated water treated mice, as described in methods, at day 24. 5-25 mice per group. P value calculated by t-test. Graphs show mean ± SD. * P<0.05. k. Lung pathology (fibrosis score) was assessed in a blinded manner, as described in methods. 5-25 mice per group. P value calculated by t-test. Graphs show mean ± SD. * P<0.05. l. Lung pathology (inflammation score) was assessed in a blinded manner, as described in methods. 5-25 mice per group. P value calculated by t-test. Graphs show mean ± SD. * P<0.05. m. WT C57BL/6J mice were given intratracheal saline (PBS) or bleomycin (BLM) on day 0, 2 and 4, as described in methods. Mice were given anti-OSM mAb (500ug/mouse every 3 days, from day −1). Body weight was monitored weekly. n. Survival was monitored and mice with >25% weight loss were euthanized. o. Total hydroxyproline (Total OHP, ug/half lung) in lung tissue was measured at day 24. 5-25 mice per group. P value calculated by t-test. Graphs show mean ± SD. * P<0.05. p. ‘New’ hydroxyproline (New OHP, ug/half lung) in lung tissue was measured in deuterated water treated mice, as described in methods, at day 24. 5-25 mice per group. P value calculated by t-test. Graphs show mean ± SD. * P<0.05. q. Lung tissue was recovered from mice at day 14 post BLM. RNA was extracted and used for RNA sequencing. Disease-relevant, tissue remodeling gene expression are shown for anti-OSM treated and isotype control mice. Graphs show individual mice with crossbars indicating mean ± standard deviation. *** FDR P<0.0005. r. Disease-relevant, fibrosis-related genes are shown for anti-OSM treated and isotype control mice. Graphs show individual mice with mean with crossbars indicating mean ± standard deviation. *** FDR P<0.0005. s. Top Ingenuity pathway analyses (IPA) are shown for isotype and anti-OSM treated mice following BLM.

### Combined IL-6 and OSM antagonism significantly reduces lung injury, inflammation, and fibrosis

Collectively, data presented above identify an inflammatory IL-6-dependent CD64^+^ myeloid activating pathway and an OSM-driven fibrotic pathway contributing to BLM-induced lung injury, inflammation, and fibrosis. We therefore hypothesized that combined blockade of IL-6 and OSM would provide both anti-inflammatory and anti-fibrotic functions, respectively, to mice exposed to BLM. To test this hypothesis, we treated mice with an anti-IL-6R mAb (i.e., the murine surrogate to TCZ), an anti-OSM mAb (as above), or a combination of both mAbs. As expected, BLM-exposed mice lost weight irrespective of mAb treatment (**Fig. 4a**), with a small proportion of mice succumbing to BLM-induced disease (**Fig. 4b**). Both anti-IL-6R or anti-OSM treatment reduced lung damage (determined by changes in TV) (**Fig. 4c**), but the combination of anti-IL-6R and anti-OSM had the greatest impact, reducing tissue volume by approximately 60% (BLM + control mAbs vs BLM + anti-IL-6R + anti-OSM, 140 ± 18.3 mm^3^ vs 60.2 ± 9.2 mm^3^). The effects of combination treatment extended to fibrotic endpoints, hydroxyproline measurements, and pathology scores, with the greatest reduction in these endpoints observed in mice given both anti-IL-6R + anti-OSM (**Fig. 4d-f**). Furthermore, using airway infiltrates as a surrogate for inflammation, anti-IL-6R treatment successfully reduced inflammation, but the greatest effect again, especially for reducing airway neutrophils, was observed in mice given both anti-IL-6R + anti-OSM (**Fig. 4g**). These data indicate that IL-6 and OSM both contribute to BLM-induced lung injury, inflammation, and fibrosis in a non-overlapping manner and that combined antagonism of these pathways provides greater benefit than either single treatment alone.

**Figure 4.**
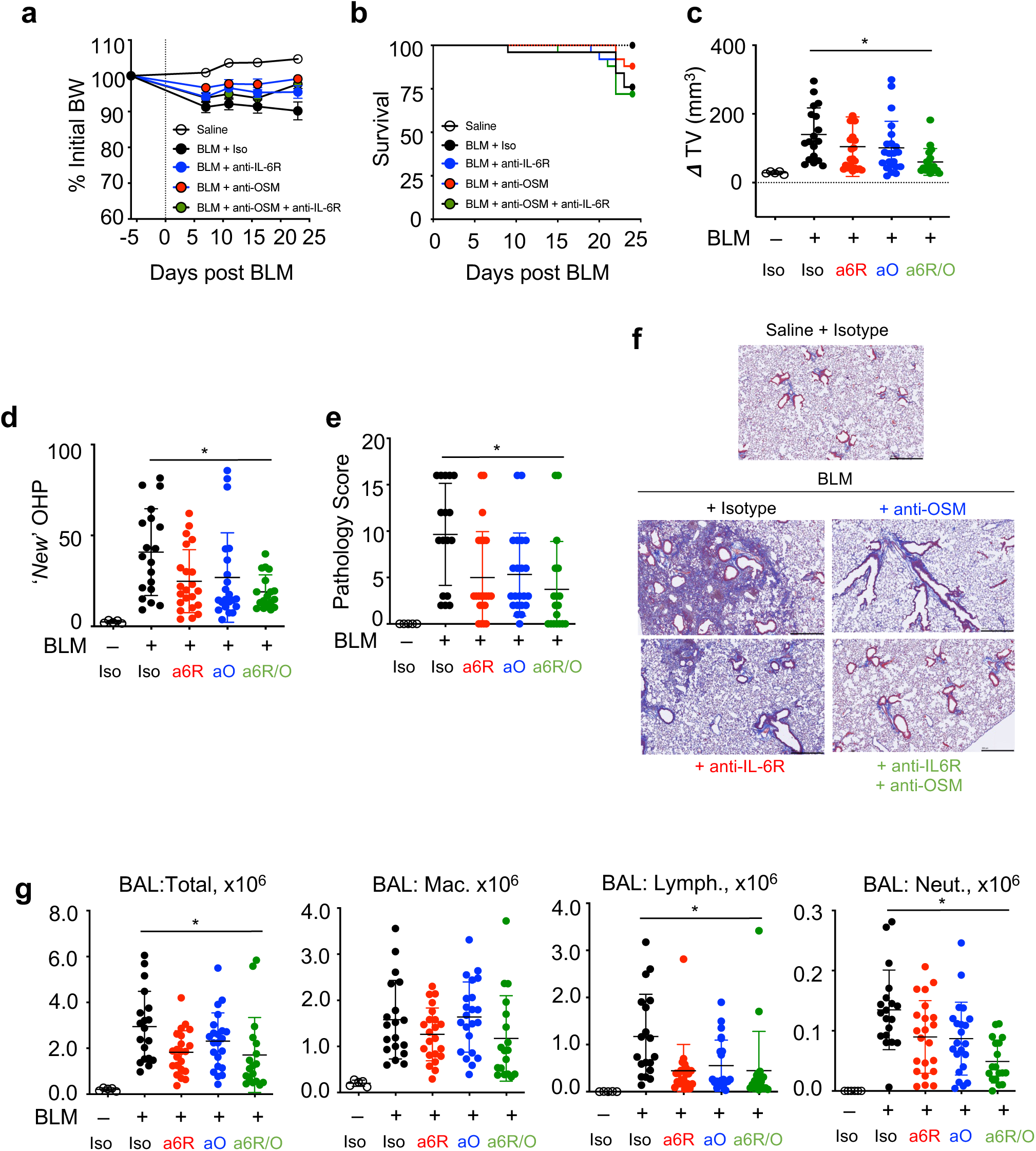
Combined IL6 and OSM antagonism significantly reduces lung injury, inflammation and fibrosis. a. WT C57BL/6J mice were given intratracheal saline (PBS) or bleomycin (BLM) on day 0, 2 and 4, as described in methods. Mice were given anti-OSM mAb + isotype, anti-IL-6R mAb + isotype, or anti-IL-6R + anti-OSMR mAb (500ug/mouse every 3 days, from day −1). Body weight was monitored weekly. 5-25 mice per group. P value calculated by t-test. Graphs show mean ± SD. * P<0.05. b. Survival was monitored and mice with >25% weight loss were euthanized. c. Tissue volume (TV) was determined at day 22, as described in methods. 5-25 mice per group. P value calculated by t-test. Graphs show mean ± SD. * P<0.05. d. ‘New’ hydroxyproline (New OHP, ug/half lung) in lung tissue was measured in deuterated water treated mice, as described in methods, at day 24. 5-25 mice per group. P value calculated by t-test. Graphs show mean ± SD. * P<0.05. e. Lung pathology (fibrosis score) was assessed in a blinded manner, as described in methods. 5-25 mice per group. P value calculated by t-test. Graphs show individual mice and mean ± SD. * P<0.05. f. Lung tissue was recovered at day 24 for sectioning and assessment of pathology. Representative masons trichrome stained sections shown. g. Bronchoalveolar lavage (BAL) was performed on mice at day 24, as described in methods. Total and differential cell counts (Macrophages, Mac; Lymphocytes, Lym; Neutrophils, Neut) were determined from each mouse. 5-25 mice per group. P value calculated by t-test. Graphs show mean ± SD. * P<0.05.

### Antagonism of IL-6R and OSM prevents myeloid cell and stromal cell activation, respectively

To identify the cell-specific transcriptional responses of BLM treated mice and how antagonism of both IL-6 and OSM pathways impact these responses, we carried out single cell RNA sequencing (scRNA-seq) on lung tissue isolated from mice given isotype control, anti-IL-6R, anti-OSM, or a combination of both anti-IL-6R and anti-OSM mAbs on day 24 post BLM. After extensive optimization of tissue dissociation, cell hashing, and the use of conventional markers, we recovered and could identify all the expected cell types in the lung (**Fig. 5a, Extended Data 6a**). We assessed the proportion of all major cell types in the lung in all treatment groups and observed only subtle shifts in the proportion of cells with or without BLM or mAb treatment. For example, there was a relative decrease in adventitial fibroblasts and increase in myofibroblasts irrespective of mAb treatment (**Extended data 6b**). Similarly, there was a reduction in AT2 airway epithelial cells and an increase in secretory cells post-BLM irrespective of mAb treatment (**Extended data 6c**). No overt changes in the proportion of endothelial cell types or lymphoid cells were observed, while a subtle increase in dendritic cells (DCs) could be seen (**Extended data 6d-6f**). This scRNA-seq approach allowed us to identify *Osmr* and *Il6ra* expressing cells, characterize their response to pathway antagonism, and help guide subsequent mechanistic studies. Firstly, *Osmr* was predominantly expressed in small airway epithelial cells (SAEC), endothelial cells (ENDO), and fibroblasts (FIB) (**Fig 5b**). In contrast, *Il6ra* was predominantly expressed in myeloid cells, as expected from data shown in **Figure 1**, but also in T cells (**Fig 5c**).

**Figure 5.**
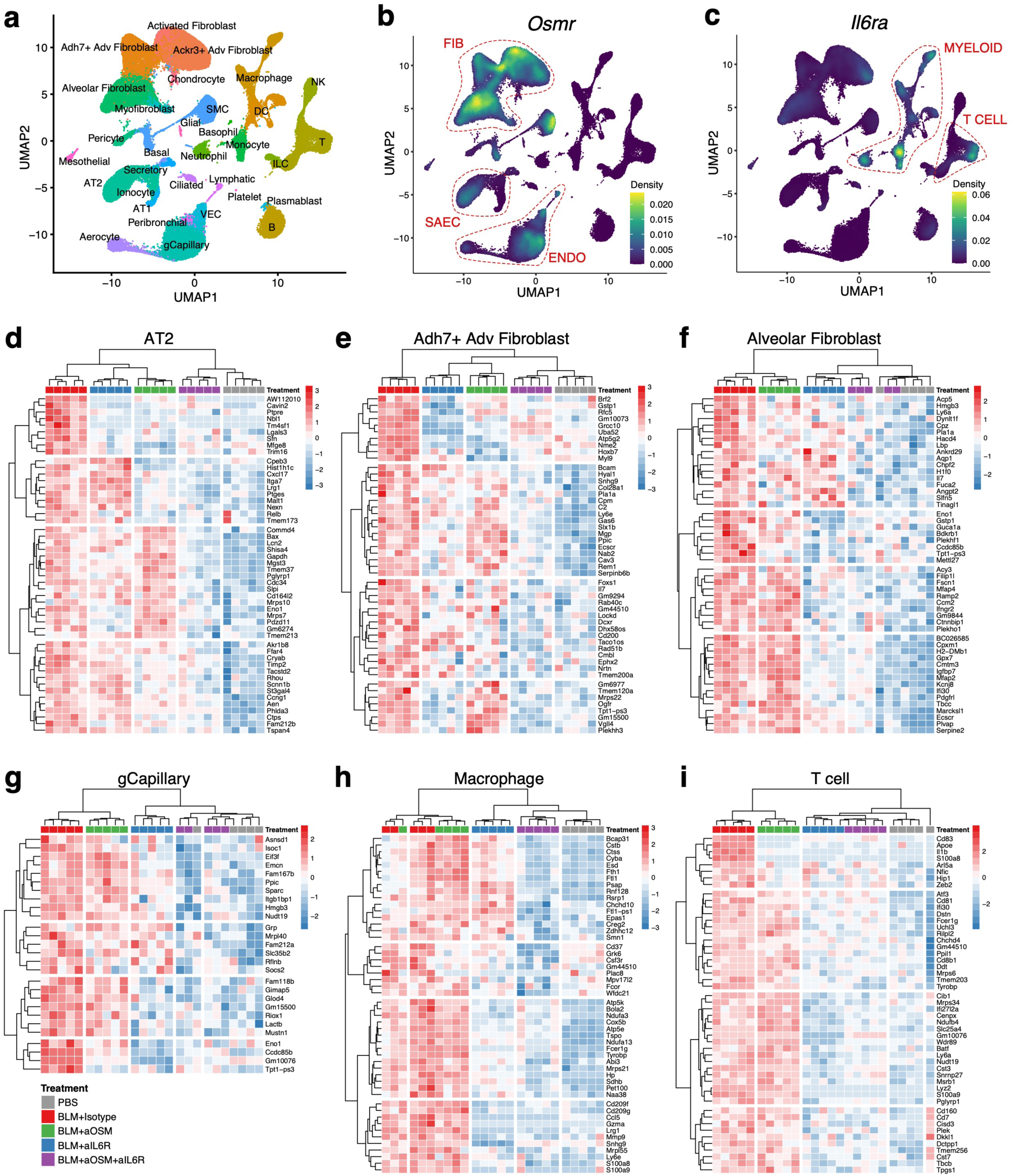
Combined IL6 and OSM antagonism has broad impact on single lung cell transcriptional responses in BLM treated mice. Lung tissue was recovered from mice at day 24 post-BLM, digested and dissociated and used for single cell RNA sequencing and analysis. a. UMAP showing cellular populations recovered and identified. b. UMAP showing *Osmr* expression density across cells. c. UMAP showing *Il6r*a expression density across cells. Hierarchical clustered heatmap showing genes (FDR <= 0.05) that were both upregulated in disease (BLM + Isotype (red treatment group, far left) vs. PBS (grey treatment group, far right) and down regulated in combination (BLM + anti-OSM + anti-IL-R6 (purple treatment group) vs. disease (BLM + Isotype (red treatment group). Individual treatment groups (anti-IL-6R (blue treatment group) and anti-OSM (green treatment group) are shown to identify impact of monotherapy. d. Genes in SAEC (AT2 cluster) shown. e. Genes in adventitial fibroblasts (Adh7+ Adv Fibroblast cluster) shown. f. Genes in alveolar fibroblasts (Alveolar Fibroblast cluster) shown. g. Genes in endothelial cells (gCapillary cluster) shown. h. Genes in macrophages (Macrophage cluster) shown. i. Genes in T cells (T cluster) shown.

To identify how combination treatment provided greater benefit than either antagonist alone, we first filtered differentially expressed genes (DEG) on FDR<0.05 that were significantly upregulated in BLM-treated mice (BLM + isotype mAbs) relative to control mice (PBS). Within these DEG, we next identified genes that were greater than 2-fold decreased in combination-treated mice (BLM + anti-IL-6R + anti-OSM), relative to the BLM-treated mice (BLM + isotype mAbs). We show in **Fig. 5d-5i** hierarchical clusters of the top 50 genes that satisfy these criteria in *Osmr*-expressing (**Fig. 5d-5g**) and *Il6ra* expressing (**Fig. 5h-5i**) cell types. This approach identified cell-specific transcriptional responses that were modulated by anti-IL-6R, anti-OSM, both, or neither, within the genes that were modulated by combination treatment (anti-IL-6R + anti-OSM). For example, within SAECs (**Fig. 5d**), BLM up-regulated the damage-associated transition progenitor (DATP) marker, *AW112010*^50^, in AT2 cells (**Fig. 5d\***) previously identified in damaged AT2 cells. Combination treatment led to >2-fold reduction in *AW112010* (a filtering criteria as described above), which could also be observed in SAECs in mice given either anti-IL-6R or anti-OSM, thus representing a common effect of antagonizing IL-6R or OSM. In contrast, combination treatment and anti-OSM alone, but not anti-IL-6R alone, prevented the SAEC expression of *Cxcl17*, a chemokine upregulated in patients with IPF^50^(**Fig. 5d\*\***), highlighting a clear and specific impact of antagonizing OSM, but not IL-6R. Similarly, combination treatment and anti-IL-6R alone effects could be observed, including prevention of *Eno1* up-regulation (**Fig. 5d\*\*\***), a serum isoenzyme biomarker associated with IPF^51^, highlighting the effect of antagonizing IL-6R, but not necessarily OSM. These cell type-specific treatment effects could be observed within adventitial and alveolar fibroblasts and endothelial cells (**Fig. 5e-g**) where *Osmr* was predominantly expressed. Interestingly, anti-IL-6R treatment also appeared to prevent the upregulation of many genes in fibroblasts, either reflecting the indirect effect of antagonizing IL-6R and/or a potential limitation of this murine study design and/or model. Although a clear effect of IL-6R antagonism could be observed across various cell types (**Fig. 5d-5i**), it was less clear to see where OSM antagonism had unique cell-specific effects. Furthermore, many genes in fibroblasts in combination-treated mice were indistinguishable from naïve mice (**Fig 5d-I**, purple and grey treatment groups), which did not appear to be impacted by either antagonist alone. These observations suggest that the transcriptional profiles observed with combination treatment is greater than the sum of each part and that the benefit of antagonizing both OSM and IL-6R is broad and impacts many cell types.

### OSM-driven pathogenic effects in human disease-relevant cells are OSMR-dependent

To better understand how OSM may contribute to human fibrotic lung disease, we first identified *OSMR* expression patterns in human IPF lung scRNA-seq data and found *OSMR* was also predominantly expressed in endothelial cells, fibroblasts, and airway epithelial cells (**Extended data 7a, 7b**), a similar pattern to that observed in mice (**Fig. 5b**). *IL6R* was also predominantly expressed in myeloid and some airway epithelial cells (**Extended data 7c**). We therefore explored the impact of OSM on primary human SAECs, endothelial cells, and fibroblasts *in vitro* (**Fig. 6a**). All three primary human cell types responded to OSM with significant STAT3 phosphorylation, whether cells were derived from healthy donors or patients with IPF (**Fig. 6b, Extended data 8a, 8b**). We next carried out a comparative transcriptional analysis of these 3 cell types following 24 hrs. of exposure to OSM (**Fig. 6c**). Of the top 10 OSM-induced transcripts in each cell type, many were commonly upregulated across all cell types (*CFI, JAK3, SOCS3, C1R, SPP1, IL1R1, CEBPD, GSDMC, and NAMPT*), with some notable cell-specific responses observed, including OSM-induced *ENNP2* (autotaxin) in SAECs, OSM-induced *IL6* in endothelial cells, and OSM-induced *S1PR1* in fibroblasts, all of which have clear roles in pulmonary fibrosis ^11,52,53^. In addition to these transcripts, several noteworthy responses were observed, including several chemokines (*CXCL12*, *CCL2*, *CXCL6*, and *CCL7*); *SERPINB4* and *MMP8* in SAECs; *LRG1*, *EGR1*, and *CXCL16* in endothelial cells; and *IGF1*, *CCR1,* and, in particular, *TNC* in fibroblasts^49^ (**Extended data 8c**). Comparative pathway analysis identified several OSM-driven inflammatory and remodeling responses across cell types, including granulocyte adhesion, epithelial-mesenchymal transition (EMT), macrophage activation, fibrosis, and wound healing, supporting a pleiotropic and broad pathogenic role for OSM (**Extended data 8d**). Finally, to identify putative OSM-induced biomarkers and pathway modulators, we compared the OSM-induced responses *in vitro* in the three cell types with observed transcriptional responses in the lung tissue of patients with IPF. OSM-driven *CCL2, CXCL1*, and *CXCL6* responses in SAECs were also observed in the IPF lung samples; OSM-driven *JAK3* and *TGFB3* in endothelial cells were observed in the IPF lung samples; and OSM-driven *CCL2*, *DIO2*, and *ADAM12* in fibroblasts were observed in the IPF lung samples (**Extended data 9a-9f**).

**Figure 6.**
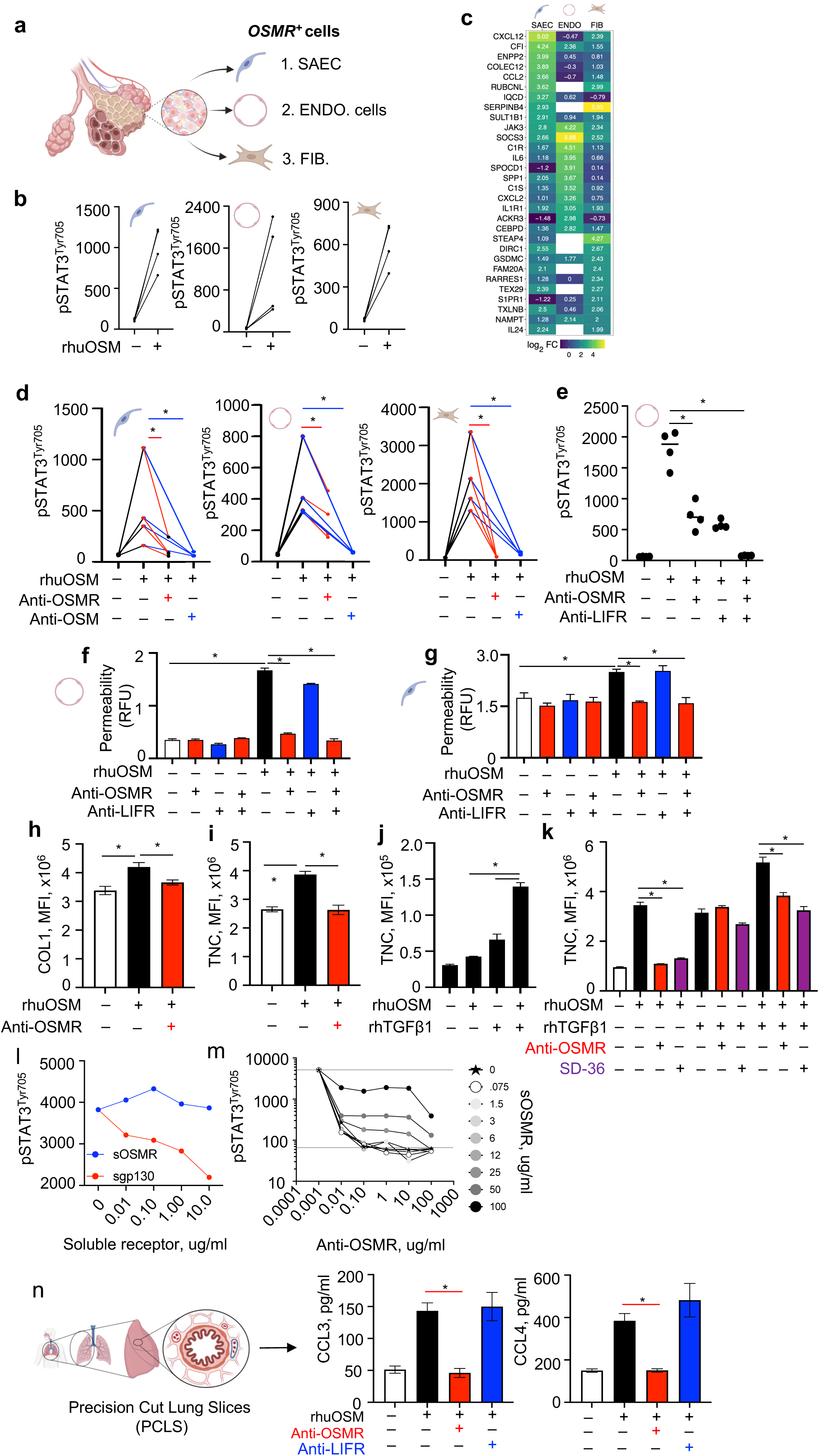
OSM mediates disease-relevant pathogenic responses in epithelial, endothelial cells and fibroblasts in an OSMR-dependent manner. a. Graphic showing OSMR-expressing shows identified from human scRNA-seq. Small airway epithelia cells (SAEC), endothelial (ENDO) cells and Fibroblasts (FIB). b. Primary human SAEC, ENDO or FIB were cultured, as described in methods, and stimulated with rhOSM (10ng/ml) for 15 mins. Cell lysates were recovered and pSTAT3^Tyr705^ was measured by MSD. c. Primary human SAEC, ENDO or FIB were cultured, as described in methods, and stimulated with rhOSM (10ng/ml) for 24hrs. RNA was extracted for RNA seq analysis. Top 10 differentially expressed genes (FDR P<0.05) are shown for each cell type. LogFC values shown in boxes. d. Primary human SAEC, ENDO or FIB were cultured, as described in methods, and stimulated with rhOSM (10ng/ml) for 15 mins. Cells were either treated with anti-OSMR (50ug/ml) or anti-OSM (10ug/ml) from 120 mins prior to OSM treatment. Cell lysates were recovered and pSTAT3^Tyr705^ was measured by MSD. P value calculated by t-test. Graphs show mean ± SD. * P<0.05. e. Primary human ENDO cells were cultured, as described in methods, and stimulated with rhOSM (10ng/ml) for 15 mins. Cells were either treated with anti-OSMR (50ug/ml) or anti-LIFR (50ug/ml) from 120 mins prior to OSM treatment. Cell lysates were recovered and pSTAT3^Tyr705^ was measured by MSD. P value calculated by t-test. Graphs show mean ± SD. * P<0.05. f. Primary human ENDO cells were cultured, as described in methods, and permeability was assessed following rhOSM (10ng/ml) treatment. Cells were either treated with anti-OSMR (50ug/ml) or anti-LIFR (50ug/ml) from 120 mins prior to OSM treatment. P value calculated by t-test. Graphs show mean ± SD. * P<0.05. g. Primary human SAEC were cultured, as described in methods, and permeability was assessed following rhOSM (10ng/ml) treatment. Cells were either treated with anti-OSMR (50ug/ml) or anti-LIFR (50ug/ml) from 120 mins prior to OSM treatment. P value calculated by t-test. Graphs show mean ± SD. * P<0.05. h. Primary human FIB cells were cultured, as described in methods, and stimulated with rhOSM (10ng/ml) for 72 hrs. Cells were treated with anti-OSMR (50ug/ml) or anti-LIFR (50ug/ml) from 120 mins prior to OSM treatment. Collagen (COL) secretion was stained and assessed using CellInsight CX7, in the scar-in-a-jar assay, as described in methods. P value calculated by t-test. Graphs show mean ± SD. * P<0.05. i. Primary human FIB cells were cultured, as described in methods, and stimulated with rhOSM (10ng/ml) for 72 hrs. Cells were treated with anti-OSMR (50ug/ml) anti-LIFR (50ug/ml) from 120 mins prior to OSM treatment. Tenascin C (TNC) secretion was stained and assessed using CellInsight CX7, in the scar-in-a-jar assay, as described in methods. P value calculated by t-test. Graphs show mean ± SD. * P<0.05. j. Primary human FIB cells were cultured, as described in methods, and stimulated with rhOSM (10ng/ml) or rhTGFβ (1ng/ml) for 72 hrs. Tenascin C (TNC) secretion was stained and assessed using CellInsight CX7, in the scar-in-a-jar assay, as described in methods. P value calculated by t-test. Graphs show mean ± SD. * P<0.05. k. Primary human FIB cells were cultured, as described in methods, and stimulated with rhOSM (10ng/ml) or rhTGFβ (1ng/ml) for 72 hrs. Cells were treated with anti-OSMR (50ug/ml) or SD-36 (10ΙM), as indicated, from 120 mins prior to OSM treatment. Tenascin C (TNC) secretion was stained and assessed using CellInsight CX7, in the scar-in-a-jar assay, as described in methods. P value calculated by t-test. Graphs show mean ± SD. * P<0.05. l. Primary human FIB cells were cultured, as described in methods, and stimulated with rhOSM (10ng/ml) for 15 mins. Cells were treated with either soluble OSMR (sOSMR) or soluble gp130 (sgp130) at indicated concentrations from 120 mins prior to OSM treatment. Cell lysates were recovered and pSTAT3^Tyr705^ was measured by MSD. m. Primary human FIB cells were cultured, as described in methods, and stimulated with rhOSM (10ng/ml) for 15 mins. Cells were treated with soluble OSMR (sOSMR) and anti-OSMR at indicated concentrations from 120 mins prior to OSM treatment. Cell lysates were recovered and pSTAT3^Tyr705^ was measured by MSD. n. Precision cut lung slices (PCLS) were prepared, as described in methods, and stimulated with rhOSM (10ng/ml) for 24hrs with anti-OSMR (50ug/ml) or anti-LIFR (50ug/ml) from 120 mins prior to OSM treatment and throughout. CCL3 and CCL4 were measured by luminex in supernatants. P value calculated by t-test. Graphs show mean ± SD. * P<0.05.

OSM binds to gp130 and heterodimerizes with either OSMR or LIFR for signal transduction. Following the observation that antagonizing the ligand OSM, but not OSMR, compromised hemoglobin and circulating platelets in patients^54,55^, it was of great interest to determine whether OSM-driven responses were mediated via OSMR and/or LIFR. We therefore tested whether antagonizing OSMR alone was sufficient to block OSM-driven responses in these 3 OSM-responsive and disease-relevant cell types.

Using an anti-human OSMR blocking mAb, we could almost completely inhibit OSM-induced pSTAT3 in SAECs and fibroblasts (**Fig. 6d**), whether derived from healthy or IPF donors (**Extended data 10a**), but could only inhibit OSM-induced pSTAT3 in endothelial cells by approximately 50% (**Fig. 6d, Extended data 10b**). The addition of anti-LIFR to anti-OSMR treated endothelial cells resulted in complete inhibition of OSM-induced pSTAT3, supporting the notion that OSM uses both OSMR and LIFR for signaling in endothelial cells (**Fig. 6e**). Collectively these data indicate that OSMR antagonism alone is sufficient to almost completely mitigate OSM-driven pSTAT3 in fibroblasts and epithelial cells, but only partially inhibit OSM-induced pSTAT3 in endothelial cells.

It has previously been reported that *in vitro* OSM can induce skin endothelial cell damage^56^. We therefore moved beyond proximal activation and signaling (pSTAT3) and assessed whether OSM also caused lung endothelial cell damage and whether this was dependent on OSMR or LIFR. Indeed, OSM induced endothelial cell disruption and permeability, compromising barrier integrity, which could be completely prevented with anti-OSMR antagonism (**Fig. 6f**). Anti-LIFR mAb treatment had little to no impact on OSM-induced permeability. Similarly, OSM-induced IL-6 and CCL2/MCP1 secretion from lung endothelial cells was largely dependent upon OSMR rather than LIFR (**Extended data 10c**). Thus, despite OSM-induced pSTAT3 being only partially mediated by OSMR, endothelial cell permeability and inflammatory cytokine production was predominantly mediated by OSMR. We carried out similar studies with primary lung SAECs grown in 3D organoids at an air liquid interface (ALI). Similar to endothelial cells, OSM could also disrupt SAEC integrity with a significant increase in permeability (**Fig. 6g**). OSM-driven SAEC permeability was also dependent upon OSMR and not LIFR, in line with pSTAT3 data (**Fig. 6d**). Taken together these data indicate that OSM can disrupt both epithelial and endothelial cell integrity, a potential pathogenic axis in ILD, and that this process is dependent upon OSMR signaling.

Most strikingly, and supporting previous studies^18^, OSM directly induced collagen (COL1) secretion from primary human fibroblasts. This process was also OSMR-dependent (**Fig. 6h**), giving some of the clearest direct mechanistic evidence for a role of OSM in human fibrotic diseases. As mentioned earlier, the ECM glycoprotein, tenascin-C (TNC), has pro-fibrotic properties^49^. In addition to OSM-induced expression of *TNC* in fibroblasts (**Extended data 8c**) and the upregulation of *TNC* in IPF lung tissue (**Extended data 9c)**, we also observed that OSM induced TNC protein secretion from fibroblasts, which was also completely dependent on OSMR (**Fig 6i**). *TNC* expression was significantly increased in skin biopsies from patients with SSc (**Extended data 10d),** and TNC protein was clearly detected in IPF lung tissue (**Extended data 10e**).

Furthermore, *TNC* expression in IPF lung tissue correlated with *OSMR* expression (**Extended data 10f**). Of note, TNC was significantly increased in mice given BLM, with anti-OSM preventing this upregulation (**Fig. 3r**), suggesting that OSM contributes significantly to TNC levels *in vitro* and *in vivo*. These data provide a second OSM-driven mechanism of fibrogenesis.

Similar to OSM, TGFβ1 has been reported to co-opt STAT3 to drive fibrotic responses^57^, induce TNC^58^, and has clear roles in a variety of fibrotic diseases^59^. It has also been reported that OSM can inhibit TGFβ1 signaling^45,60,61^. We therefore felt it was important to understand the relationship between these two pro-fibrotic mediators and determine whether TGFβ1 impacted OSM-induced TNC and COL1 secretion from fibroblasts. In contrast to previous reports^45,60,61^, we found that TGFβ1 further increased TNC and COL1 from primary lung fibroblasts when given with OSM (**Fig. 6j, Extended data 10g**) a process that was neither OSMR or STAT3-dependent (**Fig. 6k**). These data indicate that OSM- and TGFβ1-mediated fibroblast activation is independent and non-overlapping, in agreement with other studies^18^.

Finally, soluble OSMR (sOSMR) can be readily detected in circulating plasma of healthy subjects in the range of 80 ng/ml, and is significantly elevated, up to 450 ng/ml, in patients with SSc^62^. We therefore tested whether exogenous sOSMR functioned as a decoy receptor for OSM-mediated fibroblast activation and more importantly whether sOSMR interfered with anti-OSMR mediated blockade of OSM. As previously reported^63^, sOSMR alone (up to 10μg/ml) did not function as a decoy receptor and did not block OSM-induced pSTAT3 in primary human fibroblasts (**Fig. 6l**). Introduction of sOSMR only started to impact anti-OSMR mediated blockade of OSM at 25μg/ml and above, approximately 50 times what has been observed in patients with SSc^62^.

Using three disease-relevant cell types in monocultures, we demonstrate a clear role for OSMR-dependent OSM activity that drives multiple disease-relevant activation, transcriptional, and physiological responses (**Fig. 6a-m**). To determine whether OSMR is also required for OSM-driven responses in multi-cellular human lung explants, we stimulated precision cut lung slices (PCLS) with OSM and treated these cultures with anti-OSMR or anti-LIFR blocking mAbs. In agreement with the primary human monoculture systems, OSM-driven chemokine production (CCL3 and CCL4) from PCLS were dependent upon OSMR and not LIFR (**Fig. 6n**), supporting the therapeutic targeting of OSMR to prevent OSM activity in human lung diseases. Collectively these studies provide biological rationale and mechanistic data to support the therapeutic development of an OSMR antagonist to treat fibrotic lung diseases. The proposed benefit of OSMR antagonism may be further enhanced with the combined treatment with IL-6R antagonists.

## Discussion

ILDs have a devastating prognosis with life expectancy less than half a decade following diagnosis for many IPF patients. Stalling progressive fibrosis and preserving lung function are key treatment goals in patients with ILD^64^. In this study, we have characterized an IL-6-dependent inflammatory monocyte/myeloid circuit that contributes to experimental lung inflammation and pathology and correlates with several clinical observations in patients with ILD. At the heart of this axis, we observed an IL-6-driven chemokine response from myeloid cells. *In vitro*, IL-6 induced CCL18 transcription and secretion from human cells, a response which mapped to an IL-6-dependent CCL18 response in patients with ILD treated with tocilizumab. This IL-6-driven pathway has previously been implicated in, and is prognostic for, worse disease^33,34^. Similarly, in murine cells, IL-6-induced CCL2, CCL3, and CCL4 (a murine CCL18 orthologue does not exist), in agreement with previous studies showing that adenovirus-driven over-expression of IL-6 led to CCL2 secretion and lung inflammation^23^. This IL-6-driven feed-forward chemokine loop was most evident in *Il6r*^−/−^ mice treated with BLM, leading to a stunted CD64^+^ inflammatory monocyte response and reduced downstream pathology. In addition to IL-6-driven chemokine responses in myeloid cells, transcriptional analysis revealed a variety of fibrosis-associated genes in both human and murine cells (*Aqp3*, *Muc1*, *Enpp2,* and *Cd64*), in line with recent observations describing a pro-fibrotic state of macrophages ^29–31^. These broad IL-6 driven inflammatory and pro-fibrotic responses may be an important aspect of the clinical benefit with TCZ, as therapeutically targeting a single IL-6-driven chemokine, such as CCL2, has previously been unsuccessful in ILD ^65^. Although there is a lot of interest in therapeutically targeting myeloid cells, our mechanistic hypothesis suggests that dampening inflammatory monocyte activation and numbers may reduce lung function decline, as observed with TCZ, but may not necessarily impact established and progressive fibrosis.

To this end, we focused our efforts on identifying non-redundant and non-overlapping pro-fibrotic factors that could be targeted and combined with standard of care or anti-inflammatories, including IL-6R antagonists, to prevent fibrosis. Exploring the role of other gp130-dependent cytokines, we were excited to see the emerging role of IL-11 in driving fibrosis in a variety of fibrotic tissues including the liver^66,67^, lung^36,37^ and heart^38^. We also observed upregulated IL-11 in IPF and SSc tissue and in BLM-treated mice.

However, blocking the ligand with anti-IL-11 mAbs or genetically deleting the receptor did not impact inflammatory or fibrosis-related endpoints in mice treated with BLM in our hands. Similarly, in two kidney fibrosis models and a liver injury model we did not observe any impact of IL-11 blockade on fibrosis-related readouts, but instead observed a small hepato-protective effect of IL-11. Anti-IL-11 blocking mAbs could sufficiently block IL-11 activity *in vitro* and *in vivo*, had very good PK properties, and could effectively reduce HDM-induced airway inflammation, similar to previous reports^14,40–42^, so we were not overly concerned with our reagents. Although studies were carried out over multiple geographic sites using littermate mice or control mAbs^68^, there is a chance that differences in microbiota may impact the sensitivity of mice to IL-11, contributing to the discrepancies with other reports; however, we feel that this is unlikely. Finally, using primary human fibroblasts *in vitro* we found no evidence for a pro-fibrotic role for IL-11, demanding a refocus of our efforts onto other putative pro-fibrotic factors.

OSM has been very well studied in a variety of contexts, including pulmonary fibrosis^19–22,25,27,28,45–47^, but whether OSM is necessary for pulmonary fibrosis has not been reported. As we demonstrate here, using anti-OSM blocking mAbs or *Osm*^−/−^ mice, OSM has a clear role in contributing to BLM-induced lung damage and fibrosis, and this role appears to be non-redundant with the IL-6 pathway. As we previously reported^48^, microbial triggers are potent inducers of OSM from myeloid cells, and both microbial dysbiosis and myeloid cell activation have both been observed and experimentally confirmed as important contributors to pulmonary fibrosis in pre-clinical models^29–31,69,70^. Whether microbial triggers contribute to myeloid cell secretion of OSM in ILDs is unclear, but may represent a targetable upstream trigger.

The non-overlapping function of OSM with IL-6 was evident in combination studies where mice treated with both anti-OSM and anti-IL-6R mAbs received greater benefit than either anti-OSM or anti-IL-6R treatment alone. These non-overlapping functions of IL-6 and OSM have previously been implicated, with OSM-driven bronchial lymphoid structures developing independent of IL-6^71^, and with over-expression of OSM driving connective tissue fibrosis independent of IL-6^24^. To identify the cellular responses and putative mechanisms of IL-6 and OSM, we initially followed the expression of the receptors for IL-6 and OSM, which pointed us towards hematopoietic and non-hematopoietic compartments, respectively. These observations further separated these two cytokines and supported our IL-6 mechanistic studies with myeloid cells (**Fig. 1**), while also directing our OSM mechanistic studies on parenchymal and stromal cells *in vitro* and *in vivo*. Transcriptional analysis of total lung tissue and single cell analysis of OSMR-expressing cells in lung tissue from mice treated with BLM and anti-OSM highlighted a broad, but somewhat subtle single cell impact of OSM (**Fig. 3 and Fig. 5**). Tissue level analysis revealed a broad dampening of fibrotic pathways following OSM antagonism, including reduced *Mmps*, *Timp1*, a variety of collagen synthesizing genes, and *Tnc*. Single cell RNAseq analysis of lung tissue from anti-OSM treated mice focused on identifying the unique contribution of OSM antagonism in OSMR-expressing cells, relative to mice receiving both anti-IL-6R and anti-OSM. Clear transcriptional impacts in SAECs and fibroblasts were observed, and to a lesser extent in endothelial cells, while hematopoietic cells were largely unaffected (**Fig. 5**), again supporting a non-overlapping mechanism of IL-6 and OSM. Mice receiving both anti-OSM and anti-IL-6R treatment led to much more profound protection, both pathologically (**Fig. 4**) and transcriptionally in OSMR-expressing cells (**Fig. 5**), with many genes and pathways indistinguishable from non-BLM control mice.

The clearest and potentially most translatable mechanistic observations came from primary human cell studies, revealing the pleiotropic impact of dysregulated OSM in disease-relevant cells. Damage to airway epithelial and endothelial cells and subsequent activation of fibroblasts has been a long-held hypothesis contributing to the etiology and/or progression of pulmonary fibrosis^72–74^. Here we observed that OSM can contribute to all three of these processes *in vitro*, from disrupting epithelial and endothelial cell integrity and inducing inflammatory cytokine production, to promoting collagen and TNC secretion from fibroblasts. Comparative transcriptional analysis in all three cell types revealed several common pro-fibrotic transcripts upregulated by OSM (**Fig. 6**), including osteopontin (*SPP1*), which is both fibrogenic and a biomarker of progressive fibrosis^75–77^ and NAMPT, which regulates intracellular NAD and can be excreted (eNAMPT) to function as a DAMP and contribute to pulmonary fibrosis^78^. One of the most interesting OSM-driven responses was TNC, the pro-fibrotic ECM glycoprotein^49,58,79^. It has previously been reported that mice deficient in *Tnc* are protected from BLM-induced lung and skin fibrosis and resolve fibrotic scars quicker. In these studies, TNC appears to be regulated by and required for TGFβ-mediated fibrosis^49,58^. In our *in vivo* studies, *Tnc* was up-regulated in BLM-treated mice and significantly down-regulated following anti-OSM (**Fig. 3**), suggesting that OSM, in addition to TGFβ, is an important regulator of *Tnc*. Using primary human fibroblasts, TNC was secreted following OSM exposure, in an OSMR-dependent manner (**Fig. 6**). When OSM was combined with TGFβ we observed even greater TNC secretion, an OSM-driven response that was again dependent upon OSMR and STAT3. In contrast, TGFβ induced TNC was independent of OSMR or STAT3, challenging the notion that OSM inhibits TGFβ and that TGFβ-driven responses are dependent upon STAT3^57,60,61^.

In conclusion, this study has characterized an IL-6-driven mechanistic pathway and identified an important and non-redundant role for OSM in driving tissue damage and fibrosis. From *in vitro* studies, it is clear that the majority of pathogenic OSM-induced responses are dependent on OSMR, rather than LIFR, focusing our attention on the development of OSMR, rather than LIFR, antagonists, mitigating the deleterious impact of antagonizing OSM^54,55^. We hypothesize that antagonizing both OSMR and IL6R pathways will stall progressive fibrosis and preserve lung function by preventing small airway epithelial and endothelial cell damage, fibroblast activation and collagen deposition and myeloid cell activation.

## Methods

### Primary human cell cultures

Normal- or IPF-derived fibroblasts (FIB) were purchased from Lonza (#CC-2512, #CC-7231, respectively). Human pulmonary alveolar epithelial cells (SAEC) were purchased from Cell Biologics (#H-6053). Lung microvascular endothelial (ENDO) cells were purchased from Lonza (#CC-2527) and Promocell (#C-12281). All cells were thawed and expanded according to each manufacturer’s recommendation and used before or at passage 5.

### Human precision cut lung slices (PCLS)

Healthy whole lungs were received from the National Disease Research Interchange. The smallest lobe was cut free, exposing its main bronchiole, and inflated with 2% (wt/vol) low-melting-point agarose solution. Once the agarose had solidified, the lobe was sectioned. Cores of 8 mm in diameter were made in which a small airway was visible. The cores were placed in a Krumdieck tissue slicer (Alabama Research & Development Model no. MD4000), and the speed was set to produce slices at approximately 1 per 30 seconds. PCLSs (thickness, 250 μm) were transferred in sequence to wells containing Ham’s F-12 medium to identify contiguous airway segments. Suitable airways on slices were selected on the basis of the following criteria: presence of a full smooth muscle wall (cut perpendicular to direction of airway), presence of beating cilia and internal folding of epithelium to eliminate blood vessels, and presence of unshared muscle walls at airway branch points to eliminate possible counteracting contractile forces. Slices were then incubated at 37°C on a rotating platform in a humidified air/CO_2_ (95%/5%) incubator. PCLS were cultured in complete DMEM and stimulated with OSM (25ng/ml) with or without anti-human OSMR or anti-human LIFR.

### Reagents and antibodies

Recombinant Oncostatin M (OSM) and TGF-®1 was purchased from R&D systems. All cytokines in the Cytomix (TNF-α, IL-1β, IL-6 and IFN-γ) were purchased from Peprotech, reconstituted in single tube aliquots and stored at −80°C until use. Neutralizing anti-OSMR and anti-LIFR IgG1 antibodies were produced internally at Genentech. rmIL-11 and rhIL-11 were purchased from R&D and generated internally at Genentech. Anti-IL-11 mAb was used at 20mg/kg *in vivo* (MAB418, R&D or internal generated anti-mouse IL-11 mAb. sOSMR was generated internally at Genentech. SD36, the STAT3 degrader, as previously reported^77^, was purchased from MedChem Express.

### Isolation and culture of BMDMs

Femurs and tibias were harvested from mice, flushed to obtain marrow cells and passed through a 70 ΙM filter. Cells were grown on non-tissue culture treated plates in media (high glucose DMEM, 10%FBS, 10 U/ml penicillin/streptomycin, 2 mM L-Glutamine, 1 mM sodium pyruvate) containing 50 ng/ml recombinant murine M-CSF (50ng/ml, Peprotech). On day 6 macrophages were harvested, counted and re-plated for experimentation. Polarization of macrophages was carried out by addition of IL-4 (20 ng/ml, R&D Systems), IL-13 (20 ng/ml, R&D Systems), and IL-6 (20 ng/ml, R&D Systems) for 24 hours.

### Isolation and culture of human monocyte derived macrophages

Monocytes were isolated from buffy coats using RosetteSep human monocyte enrichment cocktail kit (Stem Cell, Cat No, 15028) according to manufactures instructions. Cells were grown on non-tissue culture treated plates in media (RPMI, 10%FBS, 10 U/ml penicillin/streptomycin, 2 mM L-Glutamine) containing 50 ng/ml recombinant murine M-CSF (50ng/ml, Peprotech). On day 10 macrophages were harvested, counted and re-plated for experimentation. Polarization of macrophages was carried out using IL-4 (20 ng/ml, R&D Systems), IL-13 (20 ng/ml, R&D Systems) and IL-6 (20 ng/ml, R&D Systems).

### Mice

All experiments were conducted in accordance with protocols approved by the Institutional Animal Care and Use Committee at Genentech (IACUC) and at Chugai, Singapore. WT Male (C57BL/6J), 15-20 weeks of age were purchased from the Jackson Laboratory or bred internally. The mice were kept on a 12h light-dark cycle with free access to food and water and were used in accordance with the *Guide for the Care and Use of Laboratory Animals.* IL-6 receptor knockout (*Il6r*^−/−^) mice were generated as previously described^80^. Briefly, mice with a conditional allele (*Il6r^f^*^/fl^)^79^ were interbred with Rosa26.Cre (Taconic) and maintained on C57BL/6 strain. C57BL/6 *Osm^−/−^* mice were purchased from MMRRC (C57BL/6N-*A^tm1Brd^ Osm^tm1b(KOMP)Wtsi^*/JMmucd; MMRRC Stock number 048921-UCD).

### Airway disease models

#### Mouse model of bleomycin (BLM)-induced lung injury, inflammation and fibrosis

Mice were randomized based on pre-study weights to minimize variance between experimental and control groups. For disease induction, mice were lightly anesthetized with isoflurane in an induction chamber. Once anesthetized, the animals were removed from the chamber, manually restrained, the mouth of the animal was opened, the tongue set aside and a 200uL microtip placed on the trachea. A solution of bleomycin (0.75 U kg^-1^ (DNC# 0703-3155-01; TEVA) prepared in PBS or saline was then instilled in the trachea. After delivery, animals were monitored continuously until fully awake and ambulatory. Bleomycin or saline control was administered equally in subtherapeutic doses over 3 separate days (M, W, F). Unless otherwise stated, mice were analyzed at day 24 after the 1^st^ bleomycin administration. **House Dust mite (HDM) model.** For house dust mite (HDM)-induced airway inflammation, mice were anesthetized with an i.p. injection of Ketamine 40mg/kg with Dexmedetomidine 1mg/kg. Once anesthetized, mice were treated with HDM (Greer, 10µg in 25µl) by the intratracheal route. Mice were treated with HDM on day 0, 2, 4, 14, 16, 27 and 29 and analyzed on day 30. In some experiments mice were treated with rmIL-11 (10µg in 25µl) on day 27 and 29 and analyzed on day 30. In some experiments mice were treated with anti-mouse IL-11 (500ug/mouse) on day 16, 27 and 29 and analyzed on day 30. **Broncho-alveolar lavage (BAL)** was collected in two sequential recoveries. Firstly, 500µl of ice-cold PBS was used to lavage the airspaces for cellular and analyte analysis. A second lavage of 2 x 500µl was used for additional cellular recovery. Total BAL cells and differential cell counts were determined from two BAL washes. Firstly, with 500µl of sterile cold PBS for analyte analysis and cellular recovery and then with 1ml of sterile cold PBS for cellular recovery.

### Lung cell isolation for FACS and tissue bulk RNA sequencing

Lungs were harvested, disrupted using Gentle MACS (Miltenyi) then incubated in Liberase TM, 100 ug/ml (Roche) and DNase I, 0.33 mg/ml (Roche) at 37°C for 30 minutes with gentle agitation before a final disruption using Gentle MACS (Miltenyi). The tissue was then passed through a 70-μm nylon filter to obtain a single cell suspension. Cells were treated with Ammonium-Chloride-Potassium (ACK) lysis buffer to remove erythrocytes then cells were processed for RNA analysis and FACS staining. **FACS staining and sorting.** Single cell suspensions were first treated with mouse FcR blocking reagent (Miltenyi, 130-092-575) for 10 minutes then stained for 40 minutes with combinations of antibodies to CD45 (clone 30-F11, BD Pharmingen), CD11c (clone N418, Biolegend), Siglec F (clone E50-2440, BD Pharmingen), MHCII (clone M5/114.15.2, eBioscience), CD11b (clone M1/70, Biolegend), CD64 (clone X54-5/7.1, Biolegend) and F4/80 (clone BM8, Biolegend), CD206 (clone MMR, Biolegend). Dead cells were excluded using the live/dead fixable aqua dead cell stain kit (Life Technologies, L34957). **FACS sorting.** Myeloid cells were enriched from single cell suspensions of mouse lung tissue using CD11c microbeads (Miltenyi, 130-097-059) as per manufacturer’s instructions and stained for FACS markers as described in each experiment.

### Lung histology staining and analysis

Mouse lungs were inflated with 1ml of 10% paraformaldehyde/PBS solution, embedded as a whole in paraffin and processed to 2 (5-Ιm thick) slides per animal, one series stained with Masson’s Trichrome and the other with H&E. The extent of pulmonary fibrosis was scored on the Masson’s Trichrome-stained slides according to the following criteria: (A) Interstitial fibrosis pattern—number of foci: 0, none detected; 1, ≤10; 2, ≤15; 3, >15/all sections, but distinct; 4, multifocally coalescent or locally extensive; 5, diffuse. (B) Interstitial fibrosis—size of foci: 0, none detected; 1, largest focus ≤area of ∼2 alveolar spaces; 2, largest focus ≤ area of ∼4 alveolar spaces; 3, coalescent (>4 patent alveolar spaces); 4, locally extensive (60–90% of an entire lobe); 5, diffuse (>90% of an entire lobe). (C) Total scores: number of foci × size of foci. The extent of lung inflammation was scored on the H&E-stained slides according to the following criteria: 0, normal lung (no inflammatory infiltrate); 1, minimal disease (infrequent sparsely scattered inflammatory cells); 2, mild (light perivenous/subpleural involvement); 3, moderate (many foci of small mononuclear cells extending over 3-5 alveoli); and 4, severe (generalized accumulations of small mononuclear cells involving >5 alveolar spaces).

### Hydroxyproline measurement

Total hydroxyproline was analyzed as previously described^81^ and mass spectrometry analysis was performed by Metabolic Solutions. Deuterated water labeling was used to assess new collagen synthesis. In brief, mice were injected with deuterated water (DLM-4-99.8-1000; Cambridge Isotope Laboratories) intraperitoneally two weeks prior to the end of the study at 35 mL/kg in 2 divided doses 4 h apart. Afterward, 8% deuterated water in drinking water was provided ad libitum in water bottle until the end of the study. Mass spectrometry and analysis were performed by Metabolic Solutions.

### Kidney and Liver Fibrosis models

#### Unilateral Ureteral Obstruction (UUO) model

C57BL/6NTac male mice of 6 weeks were purchased from Invivos Pte Ltd (Singapore) and were acclimated for 1 weeks before the start of study. Mice were randomized to the sham group (n=4), UUO group (n=8), and anti-IL11 antibody (Clone 5A6) treatment group (n=8). UUO surgery was operated under isoflurane anesthetized condition as previously described^82^. All monoclonal antibodies were administered at 50 mg/kg by intravenous injection one day before the surgical operation. Mice were ethically sacrificed 7 days after UUO establishment. The hydroxyproline contents in kidney was measured using hydroxyproline assay kit (BioVision). Total RNA was extracted using RNeasy Mini Kit (Qiagen), and cDNA was synthesized using a Transcriptor Universal cDNA Master (Roche Diagnostics). Quantitative PCR was performed with a TaqMan Gene Expression Assay (Thermo Fisher Scientific) and expression levels and mouse mitochondrial ribosomal protein L19 (MRPL19) was used as the endogenous reference for each sample. Relative mRNA expression values were calculated using double delta Ct analysis. **Choline-deficient, L-amino acid-defined, high-fat diet (CDA-HFD) model** C57BL/6NTac male mice of 6 weeks of age were purchased from Invivos Pte Ltd (Singapore) and were acclimated for 1 week before the start of treatments. Choline-deficient, L-amino acid-defined, high-fat diet (CDAHFD; #A06071302), was purchased from Research Diets (New Brunswick, NJ, USA) and fed to induce disease. 5P75 was fed as a normal diet. In some experiments mice were treated with anti-mouse IL-11 antibody (15 and 50mg/kg) on day 0 and 7 and analyzed on day 14. In some experiments mice were treated with rmIL-11 (R&D systems, 25 and 100ug/kg) from day 7 to 11 and analyzed on day 11. Livers were harvested and weighted. Plasma AST and ALT was measured by ALT or AST Activity Colorimetric/Fluorometric Assay Kit (Bio Vision). **Murine Nephrotoxic Nephritis model.** For nephrotoxic serum (NTS)-induced kidney fibrosis, mice were injected i.v. with 5 µl/g saline (control) or sheep anti-rat glomeruli serum (NTS; Probetex) on day 0 and monitored to 42 days post injection when they were euthanized and one kidney was submitted for histologic evaluation.

Glomerulonephritis and fibrosis were evaluated according to the following metrics. Glomerular Scoring Matrix: (0) Within normal limits. (1) Mild, segmental mesangial expansion with or without increased cellularity. (2) Moderate, segmental mesangial expansion frequently associated with increased cellularity. (3) Global membranoproliferative glomerulonephritis. (4) Glomerulosclerosis. Twenty glomeruli were scored per mouse. Fibrosis Scoring Matrix: (0) No appreciable increase in peri-glomerular fibrosis. (1) 5 or fewer individual tubules or glomeruli with surrounding fibrosis. (2) >5 individual tubules or glomeruli with fibrosis and/or 5 of few foci of interdigitating interstitial fibrosis. (3) >5 interdigitating regions of interstitial fibrosis or 1-2 regions of expansive fibrosis. (4) 3-5 foci of regionally expansive fibrosis. (5) >5 of regionally expansive fibrosis.

### Scar-in-a-jar assay

Mature extracellular collagen and tenascin-C fiber deposition was assayed as previously described^83^. Briefly, Normal- or IPF-derived human lung fibroblasts were seeded at 10,000 cells/well in complete FGM-2 media overnight. Following 2 hours pre-treatment with anti-OSMR IgG1 antibodies, isotype controls or the STAT3 degrader (SD-36), cells were incubated with OSM (10ng/mL) and/or TGF-®1 (1ng/mL) in Ficoll media for 72 hours. At harvest, cells were fixed with 4% paraformaldehyde supplemented with 4% sucrose, blocked with PBS-2% BSA and stained overnight with mouse anti-collagen 1 (#C2456; Sigma) or anti-tenascin-C antibodies (#ab3970; Abcam). The following day, cells were stained with AlexaFluor 488-conjugated anti-mouse antibodies (#A-11001; Life Technologies) and Hoechst 33342 (#H3570; Fisher Scientific) and data was acquired on the high-content imaging platform (CellInsight CX7; Thermo Fisher). All intermediate washes were performed with PBS-0.05% Tween 20.

### Cell permeability (XPerT) assay

Lung microvascular endothelial cells were seeded on gelatin-biotin-coated black plates at 30,000 cells per well and left in EGM-2MV media (#CC-3202; Lonza) to form a homogeneous monolayer for 72 hours. Where indicated, cells were pre-treated with anti-OSMR and/or anti-LIFR antibodies for 2 hours. Insult was induced by addition of OSM or Cytomix [1X = TNF-α, 0.25ng/mL; IL-1β, 0.2ng/mL; IL-6, 0.25ng/mL; IFN-γ, 1ng/mL], or PBS for controls, and plates were incubated for 20-24h. At harvest, plates were stained with Neutravidin-Dylight 488 (#22832; Thermo Scientific), washed and relative fluorescence intensity was acquired on a Synergy HT (BioTek) microplate reader. SAEC were seeded at 12,500 cells/well in Small Airway Epithelial Cell Growth Medium (SAGM) (#CC-3118; Lonza) into 96-well transwell inserts. The next day, both the apical and basolateral media was exchanged with SAGM media supplemented with 100 nM Dexamethasone (#D4902; Sigma), 5 ng/mL Human FGF-7 (#PHG0094; Thermo Fisher Scientific, 50 uM 8-Br-cAMP (#B7880; Sigma) and 25 uM IBMX (#I7018; Sigma).

Two days later, the cells were airlifted by removing the media from the apical chamber of the transwell and the media in the basolateral chamber was exchanged with DMEM:F12 containing 2 mM Glutamax, 100U/mL Penicillin, 100 ug/mL Streptomycin, and 250 ng/mL amphotericin B. The media in the basolateral chamber was change every 2-3 days until the end of the airlifting period (11-13 days). When indicated, airlifted SAEC were pre-treated with anti-OSMR and/or anti-LIFR antibodies for 2 hours. Insult was induced by addition of 12.5-25 ng/mL OSM or Cytomix [1.5-5X= TNF-α at 0.3-0.6ng/mL; IL-1β at 0.3-0.5ng/mL; IL-6 at 0.3-0.5ng/mL; IFN-γ at 1.3-1.9 ng/mL], or media for controls. After 20-24h incubation, the transwells were moved to receiver plates containing 200 uL of Transport Buffer (HBSS, 10 mM HEPES, pH 7.4) in the basolateral chamber. The apical volume was aspirated and replaced with 100 uM Lucifer Yellow (#L453; Thermo Fisher Scientific) in Transport Buffer and the quantitation of Lucifer Yellow in the basolateral chamber was measured (Ex428, Em536) after 3 hours of translocation time. The remaining Lucifer Yellow solution in the apical chamber was then aspirated, replaced with Cell Titer Glo Reagent (#G7570; Promega) and luminescence was recorded after 10 min.

### Luminex Assay

Human and murine chemokine and cytokine concentrations were assayed by Luminex technology using Bio-Plex Pro (Bio-Rad Laboratories) according to manufacturer’s instructions. Fluorescence Intensities (FI) from the labelled beads were read using FlexMaps instrument (Luminex Corp). FI from diluted standards were used to construct standard curves using Bio-Plex Manager software (Bio-Rad Laboratories) using either 4-pl or 5-pl regression type. Data is presented as average of duplicate measurements.

### Surface plasmon resonance

The binding kinetics of anti-IL11 antibody MAB418 was measured using surface plasmon resonance (SPR) on a Biacore 8k+ instrument (Cytiva). MAB418 was directly immobilized on a CM5 Series S sensorchip using amine coupling. Antibody binding was measured to mouse IL11 using a concentration series starting with 300 nM with one to three dilutions. Sensorgrams for binding of cytokine were recorded using an injection time of 2 minutes with a flow rate of 30 ul/min, at a temperature of 25°C, and with a running buffer of 10mM HEPES, pH 7.4, 150 mM NaCl, 3 mM EDTA, and 0.005% Tween 20. After injection, disassociation of the cytokine from the antibody was monitored, and the surface was regenerated between binding cycles with 10 mM Glycine HCl pH 1.7. After subtraction of a blank which contained running buffer only, sensorgrams observed for cytokine binding to MAB418 were analyzed using a 1:1 Langmuir binding model with software supplied by the manufacturer.

### MSD Assay

Murine and Human phospho-STAT3 (Tyr705) (#K150SVD-4; Meso Scale Discovery) was measured in total cell lysates per manufacturer’s recommendation.

### Immunohistochemistry (IHC)

Cells were fixed with 4% formaldehyde in PBS for 15 minutes, washed 3 times for 5 minutes in PBS, then blocked and permeabilized with a 1× PBS/5% normal goat serum/0.3% Triton X-100 solution for 1 hour at room temperature. A mouse monoclonal antibody to α-Smooth Muscle Actin (MAB1420, R&D Systems) was diluted to 15ug/ml in a 1× PBS/1% BSA/0.3% Triton X-100 solution and incubated overnight at 4°C. Slides were washed 3 times with PBS and incubated in secondary antibody at a concentration of 1ug/ml in PBS (A-11001, Thermo Fisher Scientific) for 1 hour at room temperature. Samples were washed 2 times for 5 minutes with PBS, and mounted with DAPI-containing ProLong Gold Antifade Mountant (Thermo Fisher Scientific). Images were taken on a Zeiss Axio Imager M2 microscope. A panel of 174 rabbit-derived monoclonal antibodies raised against human OSMR were screened and selected based on their specificity and sensitivity in pilot immunohistochemistry assays on OSMR-expressing formalin-fixed, paraffin-embedded cultured cell pellets. Two unique and non-competitive clones (mAbs 604 and 605) were selected for further assay optimization and staining on human tissues; both mAbs showed similar staining patterns on tissue after assay optimization, so only mAb 605 staining is presented. The optimized assay was performed on the Ultra Discovery Platform (Roche Diagnostics) using CC1 standard antigen retrieval, Discovery Antibody Block, mAb 605 diluted 3%BSA/20% human serum, OmniMap anti-Rabbit-HRP, and DAB detection. Anti-Tenascin C (TNC) staining was optimized on TNC-expressing formalin-fixed, paraffin-embedded cultured cell pellets and human skin and glioblastoma tissues; the optimized assay was performed on the Ultra Discovery Platform (Roche Diagnostics) using Protease 2 antigen retrieval, mAb DB7 (Abcam) diluted 3%BSA, OmniMap anti-Mouse-HRP, and DAB detection. Human tissues used for OSMR staining were approved by the University of California, San Francisco Institutional Review Board and adhered to the principles of the Declaration of Helsinki; written informed consent was obtained from all subjects. Fibrotic lung tissues were obtained at the time of lung transplantation from patients with a diagnosis of usual interstitial pneumonia and normal lung tissues were obtained from lungs rejected for transplantation by the Northern California Transplant Donor Network. For TNC staining, all human tissues were acquired from commercial sources under warranty that appropriate Institutional Review Board approval and informed consent were obtained.

### Pharmacokinetic analysis of anti-mouse IL-11 study

The PK of mAb418 following single i.v. dose of 10 mg/kg were conducted at Charles River Laboratories, using naive female C57BL/6 mice, age 6 to 8 weeks. Blood samples were collected by retro-orbital bleeds and the terminal blood sample was collected by cardiac stick from each animal in each dosing group at various timepoints up to 14 days post-dose with three mice per time point and processed to collect serum. Time course of PK profiles and PK parameters are shown in **Extended data Fig2e.** Serum samples were analyzed for test article concentrations by enzyme-linked immunosorbent assay. Mouse anti-mouse IgG2a (BD Biosciences) was used as the capturing reagent and anti-mouse IgG2a specific antibody conjugated to horseradish peroxidase (HRP) (GeneTex), was used as the detection reagent. The serum concentration versus time data were used to calculate PK parameters in mouse using noncompartmental analysis (Phoenix WinNonlin, Version 6.4.0.768; Certara). Nominal sample collection times and nominal dosing solution concentrations were used in the data analysis.

### ELISA/Luminex

For chemokines were measured by Luminex using a MILLIPLEX MAP Mouse Immunoglobulin Cytokine/Chemokine Magnetic Beat Panel (cat. no. MCYTMAG-70K-PX32, Millipore).

### RNA sequencing (RNA-seq), single cell RNA-sequencing (scRNA-seq) and gene expression analysis

The concentration of total RNA samples was determined using NanoDrop 8000 (Thermo Scientific) and the integrity of RNA was determined by Fragment Analyzer (Advanced Analytical Technologies). 0.5 µg of total RNA was used as input material for library preparation using TruSeq RNA Sample Preparation Kit v2 (Illumina). Size of the libraries was confirmed using Fragment Analyzer (Advanced Analytical Technologies) and their concentration was determined by a qPCR-based method using Library quantification kit (KAPA). The libraries were multiplexed and then sequenced on a HiSeq2500 (Illumina) to generate 30M single-end 50 base pair reads. RNA was quantified using Nanodrop 8000 (Thermo Scientific) and integrity was measured using the Bioanalyzer RNA 6000 Pico Kit (Agilent). Libraries were prepared using the TruSeq RNA Library Prep Kit v2 (Illumina) with 100-500 ng of input and amplified using 10 cycles of PCR. Libraries were multiplexed and sequenced on a HiSeq 2500 System (Illumina), resulting in 15M single-end 50 bp reads per library. Alignment, feature counting, normalization, and differential expression analysis were performed similar to as described previously (14). In brief, HTSeqGenie was used to perform filtering, alignment to GRCm38, and feature counting. Normalized counts per million (CPM) values were computed as a measure of gene expression. Pairwise differential expression analysis was performed using voom and limma. For differential gene-expression analysis significant genes were filtered and identified as Benjamini-Hochberg corrected p <0.05. Pathway analysis was performed with Ingenuity Pathway Analysis (IPA) software (Qiagen) using the “Diseases and Disorders” pathway module. Heat map Euclidean clustering of genes was performed by plotting log 2-transformed fold change values for each replicate sample and each gene (log 2 floor set at −3 for heat map). Colored boxes indicate the degree of fold change. For gene expression by qRT-PCR, cells or tissues were lysed using Buffer RLT (Qiagen) and total RNA extracted using RNeasy Mini Kit with on-column DNase digestion (Qiagen) and reverse transcribed to cDNA using iScript kit (Bio-Rad). Gene expression was analyzed using TaqMan Gene Expression Assay (Applied Biosystems). *Ccl2* (Mm00441242_m1), Ccr2 (Mm99999051_gH), *Cd64* (Mm00438874), *Col1a1* (Mm00801666_g1), *Col1a2* (Mm00483888_m1), *Fn1* (Mm01256744_m1), *Tgfb1* (Mm01178820_m1), *Arg1* (Mm00475988_m1), *Cd206* (Mm01329362_m1), *Nos2* (Mm00440502) and *CCL18* (Hs00268113_m1). Target gene expression was normalized by housekeeping genes *Hprt1* (Mm01545399_m1) and *Rpl19* (Mm02601633_g1) for mouse transcripts, and by GAPDH (Hs02786624_g1) for human transcripts.

### Lung cell isolation for single-cell RNA sequencing

Lungs were harvested and processed in single cell suspensions such as described above, with modifications taken from previous protocol optimized for single cell transcriptomics^84^. After tissue digestion, RBC lysis and FcR blocking, samples were co-stained with lineage marker antibodies (CD45-APC, eBioscience #17-0451-82; CD31-FITC, Biolegend #102405; EpCAM-PE, Biolegend #118205), cell hashing antibodies (TotalSeq B 1-5, Biolegend) and with a live/dead stain (7-AAD, BD #559925). After staining, samples were normalized in total number of cells recovered, pooled per treatment group and FACS sorted to isolate immune, epithelial, endothelial and stromal cell fractions.

### Single-cell RNA sequencing

Sorted cell fractions were loaded on the Chromium Next GEM Single Cell 3’ v3.1 kit with a target of 16k cells per sample. For all samples, viabilities were greater than 80% and cell concentrations ranged between 700 and 1,600 cells/ul. Following the manufacturer’s instructions, we loaded approximately 26400 cells into each channel. Library generation was performed following the manufacturer’s protocol 3’ v3.1 with cell surface protein. Briefly, cell suspensions were encapsulated in Gel-emulsion beads followed by reverse transcription. The Gel Beads-in-emulsion (GEMs) were then broken open releasing the pre-amplification cDNA which was cleaned and amplified. cDNA was then converted into sequencing libraries via fragmentation followed by ligation of the sequencing adapters and final addition of Illumina compatible dual-index primers. Libraries were sequenced on a NovaSeq 6000 according to the manufacturer’s specifications.

### scRNA-seq data processing and analysis

scRNA-seq sequencing reads collected from the *in vivo* anti-OSM and anti-IL6R treatment experiment were filtered and aligned using cellranger v6.0.1 (10x Genomics). This data was then analyzed in the R v4.2.1 statistical computing environment using the Seurat v4.2 R package^85,86^. CMO data was normalized through CLR (centered-log ratio) transformation and HTO data was demultiplexed using k-means clustering. Only cells with ≥ 500 genes were retained for analysis, normalization was performed using the log-normalization method, and 2,000 highly-variable genes were selected for sample integration and clustering using the mean/variance regression method. Sample integration and batch correction was performed using the anchor-based sample integration workflow. Clustering was performed using 25 PCA components on a k=20 shared nearest neighbor (SNN) graph using the Louvain algorithm. UMAP dimensional reductions were performed using the uwot R package. Marker selection was performed using the Wilcoxon Rank Sum test on the integrated sample data for each cluster against all clusters. For all analyses, percent of cells expressing a given gene (detection rate) was defined as the presence of ≥ 1 UMI for the given gene. Cell type annotation was performed manually based differentially expressed genes from a one-vs-all comparison meeting the criteria of average log2 fold-change ≥ 0.5, a minimum detection rate ≥ 20%, and either detection rate ≥ 50% within a cluster or ≤ 10% within all other clusters. Withing each cell type annotations, gene counts were pseudo bulked by summing count data per gene across each cell type for each mouse. Differential gene expression was calculated using voom+limma with multiple-hypothesis correction of p-values via the Benjamini-Hochberg method. Heatmaps were generated using the pseudobulk log_2_ normalized CPM expression matrix with both rows and columns hierarchically clustered using Euclean distance and Ward’s linkage. The top 50 most variable genes that were upregulated in the BLM + Isotype vs. PBS differential expression, downregulated in the BLM+aOSM+aIL6R vs. BLM+Isotype comparison, with corrected p ≤ 0.05 and had at least a 2-fold change difference between the two differential expressions were plotted.

**All data sets are uploaded to GEO:** accession GSE263043

## Acknowledgements

We thank members of the Wilson laboratory and Immunology Discovery department, Genentech, for discussion and reading of the manuscript and editorial assistance. The authors thank Matthew Taylor (PTPK, Genentech) and Pamela Chan (BCP, Genentech) for assistance with PK analysis. We would like to thank the vet staff at Genentech, Catherine Sohn, Dorie Montoya, Raul Garcia-Gonzalez, Emmanuel Chua, Jill Yamada, Katelyn McEachin, Ryan Scott, Fidel Guardado and Jennifer Cosino for the meticulous care and support for all animal work. We thank CK Poon, Jim Cupp, Terence Ho, Qi Xuan Tan, Andres Paler Martinez, Jovencio Borneo, Cole Meyers, Kevin Le in the FACS/Luminex laboratory for their support. We thank Jingyu Diao and Ke-Jung Huang for their initial work optimizing and performing permeability assays. We thank Alison Huynh, Jessica Mills, Sean Flanagan, Shannon Hambro, Victor Nunez, and their team for the invaluable support in animal work. We thank Andres Paler Martinez for Luminex analysis, Zhiyu Huang and Alexander Arlantico for assistance with mouse experiments, Mariela Del Rio for mouse colony management. The authors thank the Next Generation Sequencing group, Genentech, for all of their assistance, guidance, and support with RNA sequencing. We apologize to our many colleagues whose primary work could not be cited owing to space constraints.

## Author Contributions

– R. A. R., E. D., H. S. D., S. H., C. J. W., R. G., G. T., D. X., conceptualized, designed, performed and analyzed a variety of experiments.
– D. R., carried out FACS staining and analysis.
– H. B.,S. J., C. L. E., A. A., A. W., S. Y., A. S., S. A., M. K. performed *in vivo* experiments.
– T. R., X. G., J. L., performed human fibroblast, epithelial and endothelial cell experiments.
– C. A., D. J. D., and P. C., performed and analyzed histology data.
– J. V. H., A. R. A., S. U., analyzed RNA-sequencing data sets.
– J. B., G. N., generated mAbs and carried out SPR studies.
– R. Y. carried out PK studies.
– R. A. P., C. K-W., W. F. J., provided PCLS.
– Y. S., J W., performed and analyzed OSM-induced pSTAT3 assays.
– J. R. A., conceptualized experiments and edited the manuscript.
– M.S.W., N. R. W., J. R. A., J. V. H., conceptualized the project, executed some experiments, wrote and edited the manuscript.

---

**Extended Data 1.**
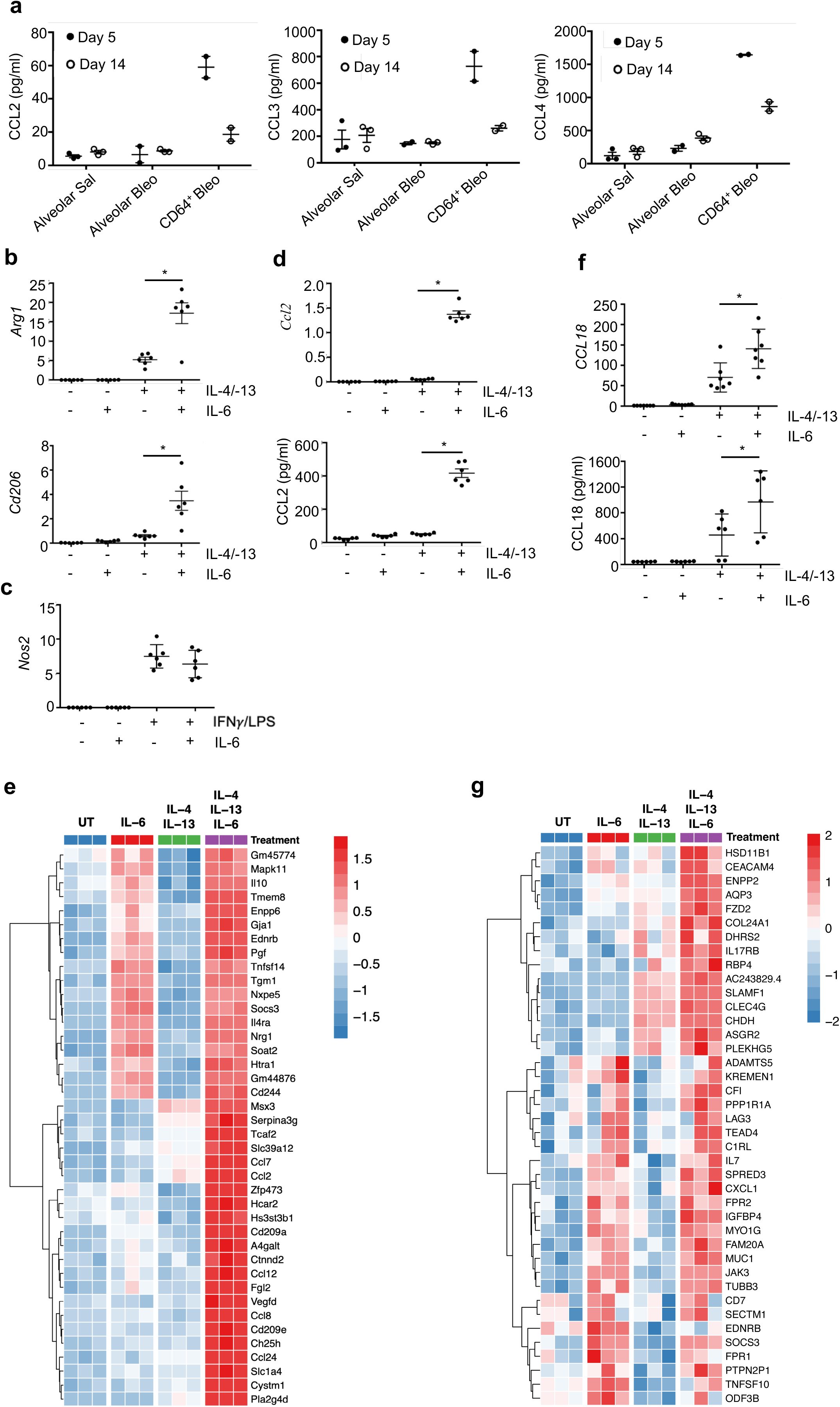
IL-6 activates myeloid cells and drives inflammatory and fibrotic programs. a. Alveolar macrophages (CD45^+^CD11c^+^SigF^+^) and CD64^+^ (CD45^+^CD11c^+^SigF^−^MHCII^+^CD11b^+^CD64^+^) macrophages were FACS sorted from the lungs of WT mice 5 and 14 days post BLM treatment. Sorted cells were then placed in culture for 24 hours and CCL2, CCL3 and CCL4 was analyzed using Luminex. Graphs show mean of two independent experiments ± SD. * P<0.05. b. WT BMDMs were treated with IL-4 and IL-13 +/– IL-6 for 24 hours before gene expression analysis was assessed by qRT-PCR (n=6). Graphs show mean ± SD. * P<0.05. c. WT BMDMs were treated with IFN© and LPS +/– IL-6 for 24 hours before gene expression analysis was assessed by qRT-PCR (n=6). Graphs show mean ± SD. * P<0.05. d. WT BMDMs were treated with IL-4 and IL-13 +/– IL-6 for 24 hours with CCL2 measured in supernatant by Luminex and gene expression assessed by qRT-PCR (n=6). Graphs show mean ± SD. * P<0.05. e. WT BMDMs were treated with IL-4 and IL-13 +/– IL-6 for 24 hours before RNA was extracted for RNA-sequencing. Heat map showing the most differentially upregulated genes between untreated and IL-4 + IL-13 +IL-6 treatment across all treatment groups (top 40 by fold change and adjusted P value <0.05, n=3). f. Monocyte derived macrophages (MDMs) were generated from healthy donors and polarized with IL-4 and IL-13 +/– IL-6 for 24 hours. CCL18 mRNA and protein was measured using qRT-PCR and ELISA, respectively (n=7). Graphs show mean ± SD. * P<0.05. g. Monocyte derived macrophages (MDMs) were generated from healthy donors and polarized with IL-4 and IL-13 +/– IL-6 for 24 hours. RNA was extracted for RNA sequencing. Heatmap showing the most differentially upregulated genes are between untreated and IL-4 + IL-13 +IL-6 treatment across all treatment groups (top 40 by fold change, adjusted p<0.05, n=3).

**Extended Data 2.**
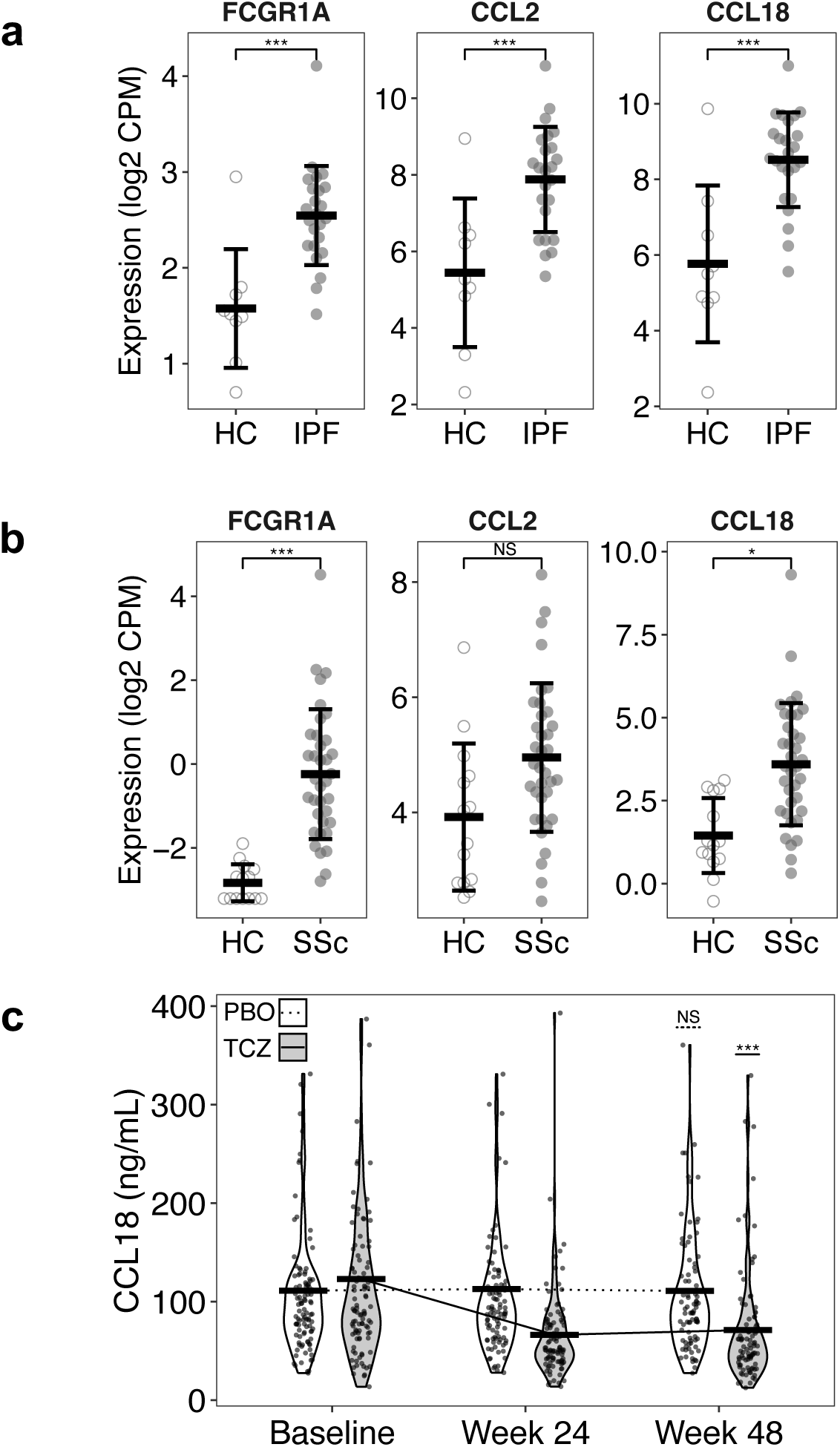
Correlates of IL-6-driven responses in human IPF lung and SSc skin and IL-6-dependnet CCL18 in serum. a. Lung biopsies were obtained from healthy controls (n=9) and IPF patients (n=24), RNA-seq was carried out and *FCGR1A (CD64), CCL2* and *CCL18* transcripts analyzed. Graphs show normalized log2 CPM, mean ± SD. ***P<0.0005, NS not significant. b. Skin biopsies obtained from healthy controls (n=14) and SSc patients (n=36) participating in the focuSSced clinical trial and *FCGR1A (CD64), CCL2* and *CCL18* transcripts measured at baseline. Graphs show normalized log2 CPM, mean ± SD. * P<0.05, ***P<0.0005, NS not significant. c. Serum was obtained from SSc patients (n=207) participating in the focuSSced clinical trial and serum CCL18 was measured at baseline and after 24 or 48 weeks of PBO (n=104) or TCZ (n=103) treatment. P values calculated by t-tests between baseline and week 48. Graphs show ng/mL and mean (bar). *** P<0.0005, NS not significant.

**Extended Data 3.**
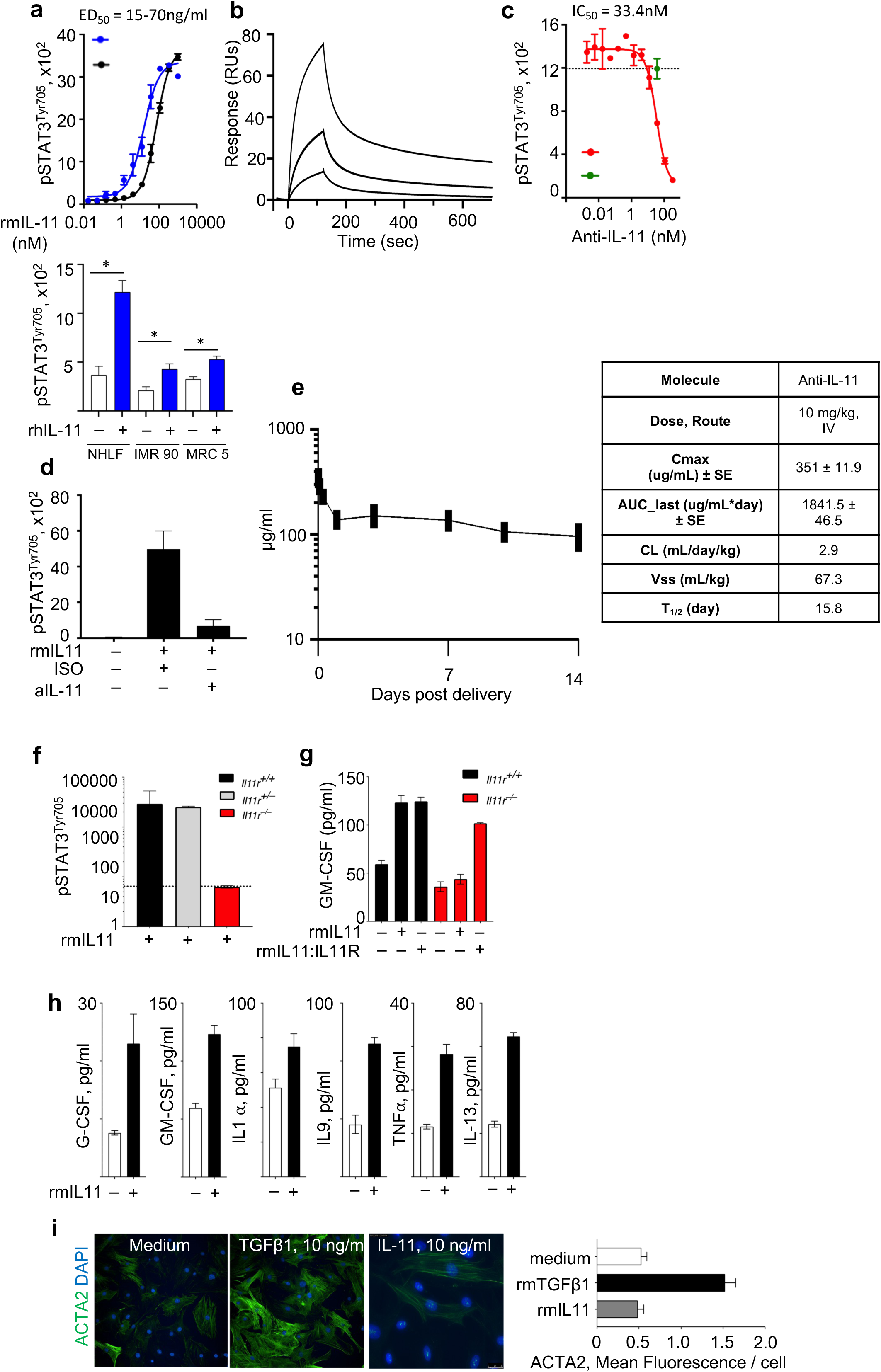
IL-11 mediated activation of STAT3 and induction of inflammatory cytokines, but not α−SMA. a. Murine LGg cells were stimulated with increasing concentrations of rmIL-11 from two different sources (murine, top, blue= R&D systems, black= in house generated, human, bottom, R&D systems). pSTAT3^Tyr705^ was measured by MSD after 15 mins. Primary human normal lung fibroblasts (NHLF) of fibroblast cell lines (IMR 90, MRC 5) were stimulated with rhIL-11 (10ng/ml). pSTAT3^Tyr705^ was measured by MSD after 15 mins. b. Binding kinetics of anti-mouse IL-11 (MAB418) determined using surface plasmon resonance (SPR). c. *In vitro* inhibition of IL-11-induced pSTAT3^Tyr705^ by anti-mouse IL-11 (MAB418, red line) or isotype rat IgG (green). IC_50_ = 33.4nM. pSTAT3^Tyr705^ was measured by MSD after 15 mins. d. *In vivo* inhibition of IL-11-induced pSTAT3^Tyr705^ by anti-mouse IL-11 (MAB418). Mice were treated with anti-IL-11 (250ug) on day 0. On day 1 mice were given 25ng of rmIL-11 via the intra tracheal route. After 15 mins, mice were euthanized and whole lungs removed. pSTAT3^Tyr705^ was measured by MSD after 15 mins in whole lung lysate. e. Pharmacokinetic (PK) properties of anti-IL-11 were characterized following a single intravenous (IV) dose of 10 mg/kg in female C57BL/6 mice. f. Primary mouse lung-derived fibroblasts were isolated from *Il11*^+/+^, *Il11*^+/–^ and *Il11*^−/−^ mice and stimulated with rm-IL-11 (10ng/ml). pSTAT3^Tyr705^ was measured by MSD after 15 mins. g. Primary mouse lung-derived fibroblasts were isolated from *Il11*^+/+^ and *Il11*^−/−^mice and stimulated with rm-IL-11 (10ng/ml) or rmIL-11 fused with IL-11R. After 24 hrs., GM-CSF secretion was measured by luminex. One of 3 experiments shown. h. Primary mouse lung-derived fibroblasts were isolated from WT mice and stimulated with rmIL-11 (10ng/ml). Cytokine and growth factor secretions were measured by Luminex after 24hrs. i. Primary mouse lung-derived fibroblasts were isolated from WT mice and stimulated with rmTGFβ (10ng/ml) or rmIL-11 (10ng/ml). ACTA2 ((SMA) was measured after 72hrs by immunofluorescence.

**Extended Data 4.**
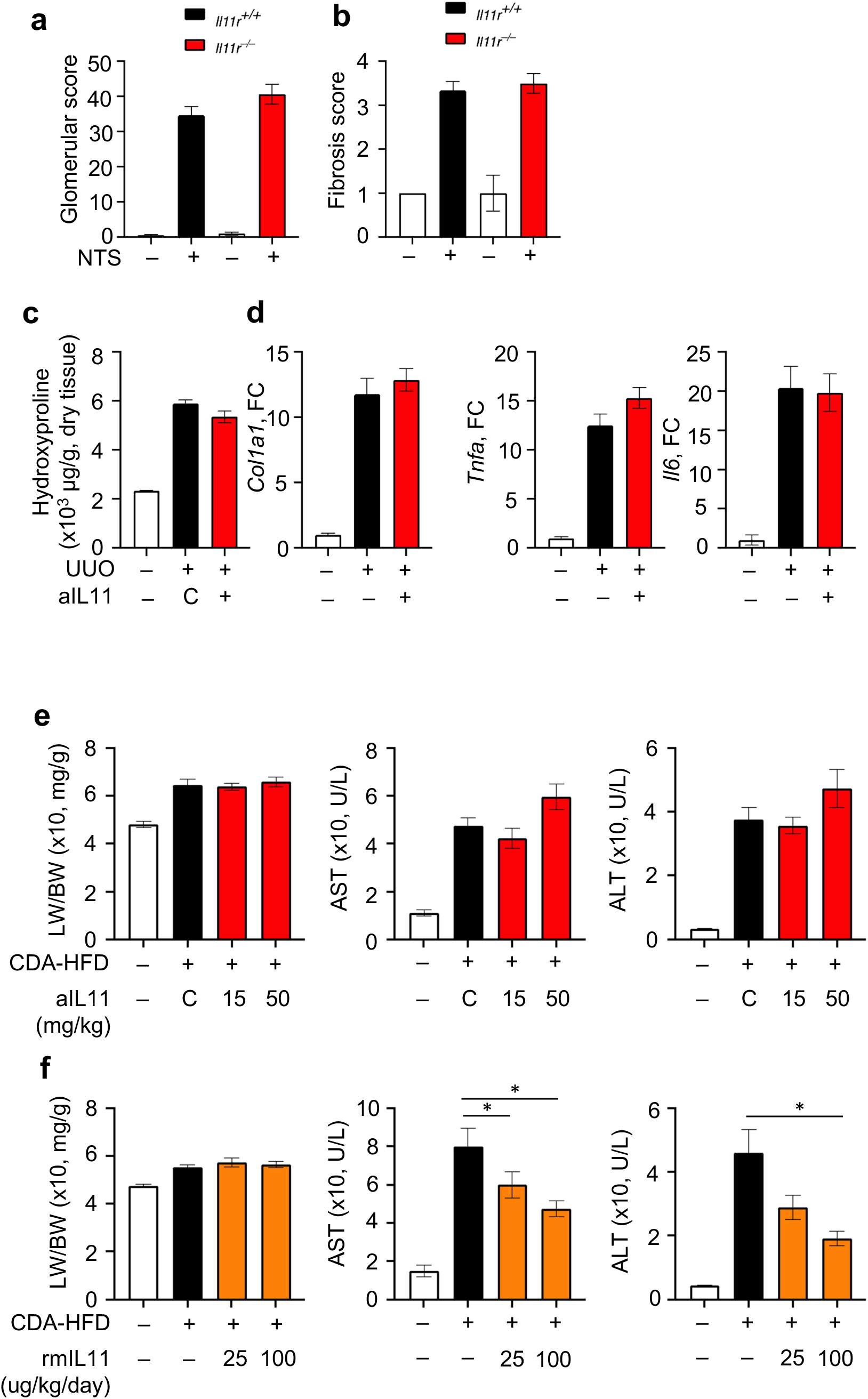
IL-11 is not required for experimental kidney fibrosis or liver damage. a. WT (*Il11r*^+/+^) and IL-11R-deficient (*Il11r*^−/−^) C57BL/6J mice were exposed to nephrotoxic serum (NTS), as described in methods. Glomerular score as described in methods, was assessed in a blinded manner on day 42. b. Kidney fibrosis, as described in methods, was assessed in a blinded manner on day 42. c. WT C57BL/6J mice underwent UUO surgery, as described in methods. Mice were given isotype control Ab (C) or 50mg/kg of anti-mouse IL-11 (aIL11, Clone 5A6) one day before surgery. Hydroxyproline (ug/g dry tissue) was assessed on day 7. d. Gene expression was measured in homogenized kidney tissue on day 7 post-surgery. e. WT C57BL/6J mice were fed a **Choline-deficient, L-amino acid-defined, high-fat diet** (CDA-HFD) *ad libitum*. 5P75 was the normal diet. Mice were treated with isotype control Ab (C), 25mg/kg or 50mg/kg of anti-mouse IL-11 (clone 5A6) on day 0 and day 7. Liver weight to body weight ratio, serum AST and serum ALT was measured on day 14. f. WT C57BL/6J mice were fed a CDA-HFD as above. Mice were given saline or rmIL-11 (25 or 100ug/kg/day, R&D) from day 7-11 and assessed at day 14.

**Extended data 5.**
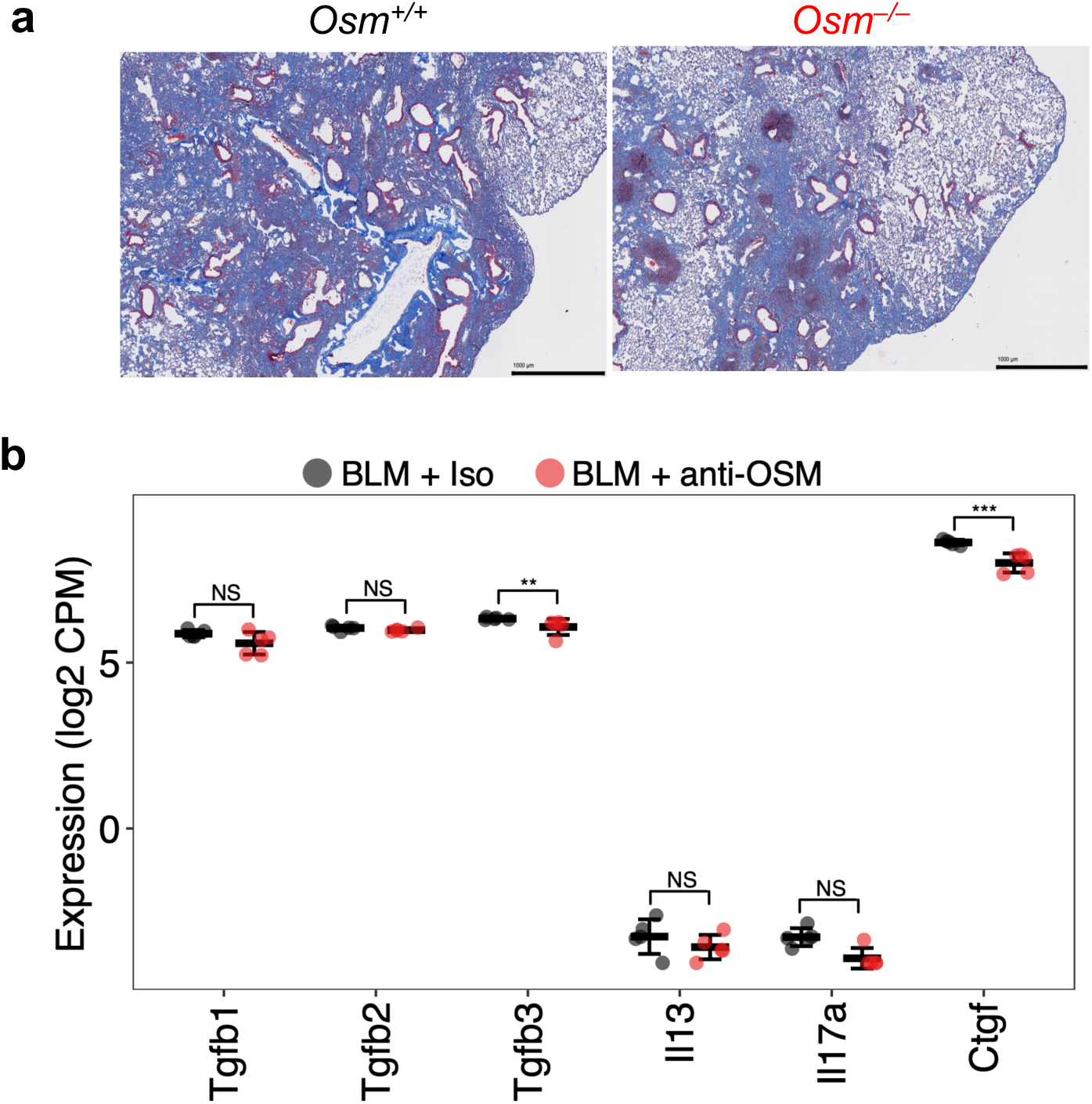
Oncostatin M (OSM) antagonism reduces tissue fibrosis, but not canonical pro-fibrotic cytokines or growth factors. a. WT (*Osm*^+/+^) and OSM-deficient (*Osm*^−/−^) C57BL/6J mice were given intratracheal saline (PBS) or bleomycin (BLM) on day 0, 2 and 4, as described in methods. Lung tissue was recovered at day 24 for sectioning and assessment of pathology. Representative masons trichrome stained sections shown. b. Lung tissue was recovered from mice at day 14 post BLM. RNA was extracted and used for RNA sequencing. Disease-relevant, pro-fibrotic cytokines and growth factor genes are shown for anti-OSM treated and isotype control mice. Graphs show individual mice with crossbar indicating mean ± standard deviation. NS not significant, ** FDR P<0.005, *** P<0.0005.

**Extended data 6.**
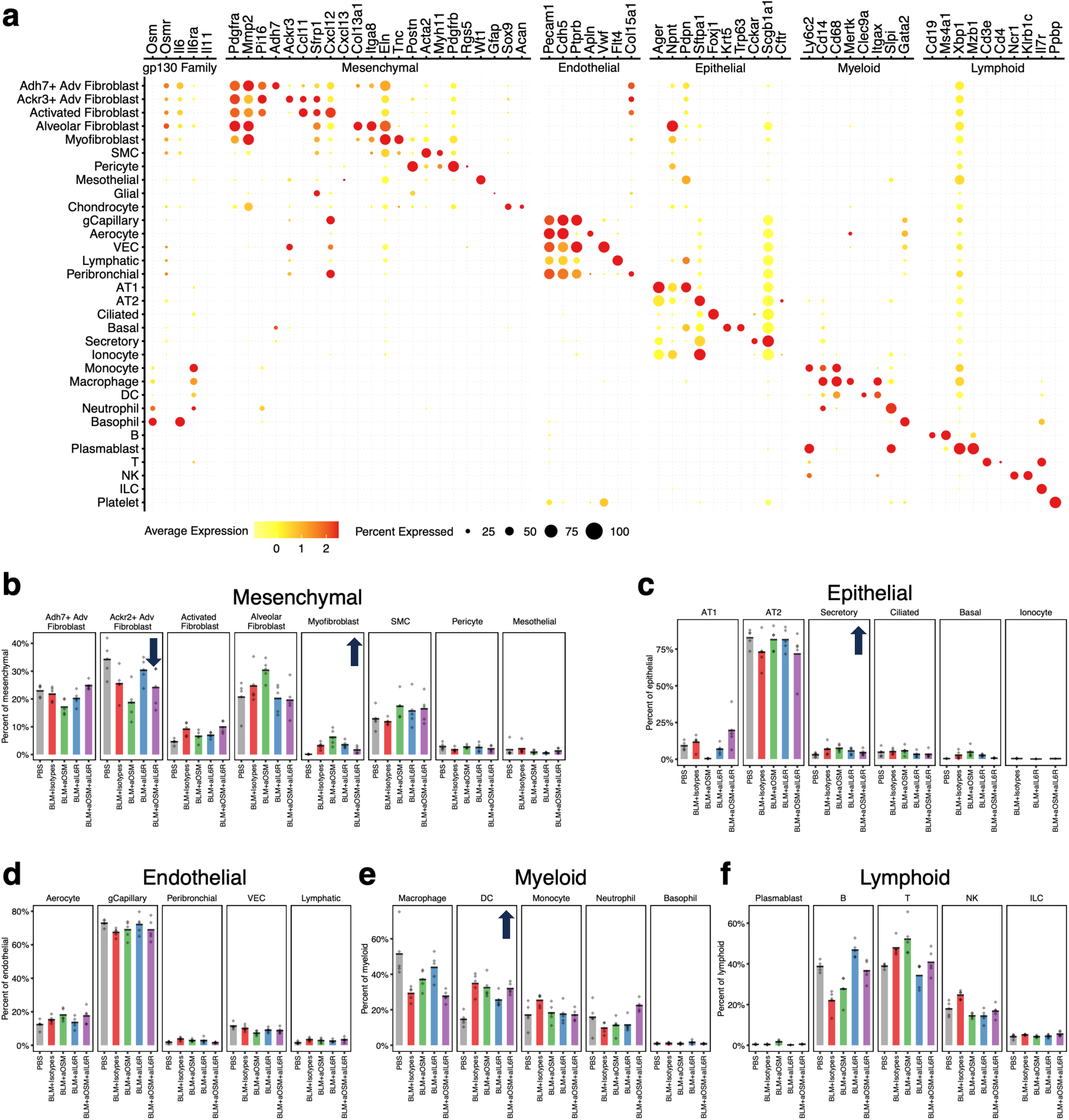
Lung cell transcriptional identities and profiles and proportional changes following BLM and mAb treatment. Lung tissue was recovered from mice at day 24 post-BLM, digested and dissociated and used for single cell RNA sequencing and analysis. a. Dot plot showing cell identities (left) and representative gene expression. Size of dot refers to percentage of cells expressing gene and color refers to average expression. b. Proportional shifts in mesenchymal cell populations following BLM and mAb treatment for individual mice (points) and mean (bar height). c. Proportional shifts in epithelial cell populations following BLM and mAb treatment for individual mice (points) and mean (bar height). d. Proportional shifts in endothelial cell populations following BLM and mAb treatment for individual mice (points) and mean (bar height). e. Proportional shifts in myeloid cell populations following BLM and mAb treatment for individual mice (points) and mean (bar height). f. Proportional shifts in lymphoid cell populations following BLM and mAb treatment for individual mice (points) and mean (bar height).

**Extended data 7.**
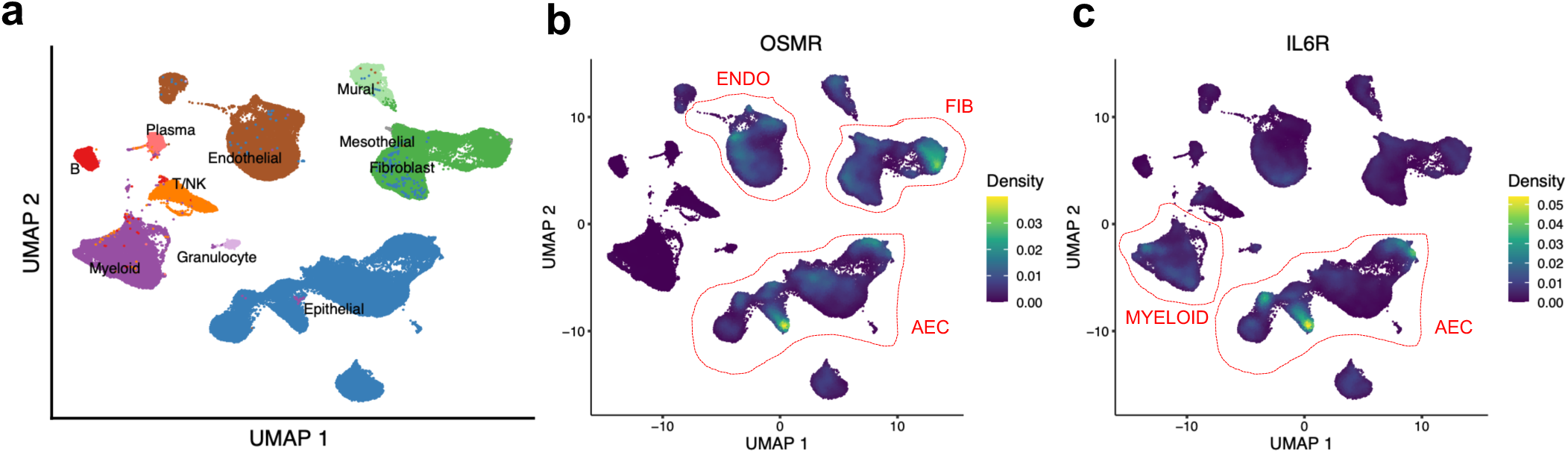
Single cell transcriptional analysis of human ILD lung tissue. a. UMAP showing cellular populations recovered from digested lung tissue. b. UMAP showing *OSMR* expression density across cells. c. UMAP showing *IL6R* expression density across cells.

**Extended data 8.**
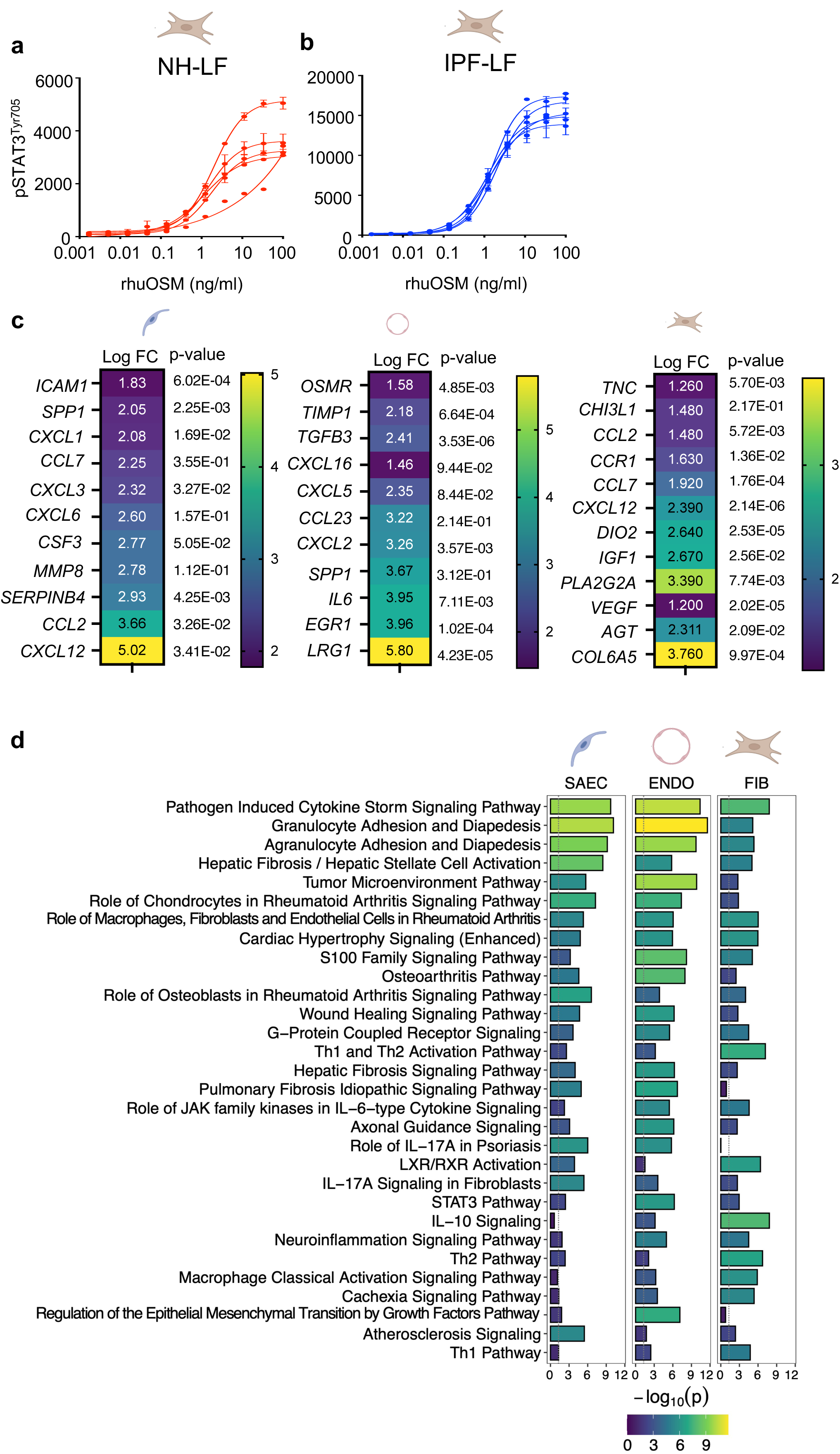
OSM-driven pathways in disease-relevant cells. a. Normal primary human lung FIB (NH-LF) were cultured, as described in methods, and stimulated with rhOSM at the indicated concentrations for 15 mins. Cell lysates were recovered and pSTAT3^Tyr705^ was measured by MSD. b. IPF-derived human lung FIB (IPF-LF) were cultured, as described in methods, and stimulated with rhOSM at the indicated concentrations for 15 mins. Cell lysates were recovered and pSTAT3^Tyr705^ was measured by MSD. c. Primary human SAEC, ENDO or FIB were cultured, as described in methods, and stimulated with rhOSM (10ng/ml) for 24hrs. RNA was extracted for RNA seq analysis. Disease-relevant differentially expressed genes (P<0.05) are shown for each cell type. Log_2_FC values shown in boxes with p-values shown to the right. d. Primary human SAEC, ENDO or FIB were cultured, as described in methods, and stimulated with rhOSM (10ng/ml) for 24hrs. Ingenuity pathways analysis (IPA) analyses of OSM-driven responses are shown for each cell type.

**Extended data 9.**
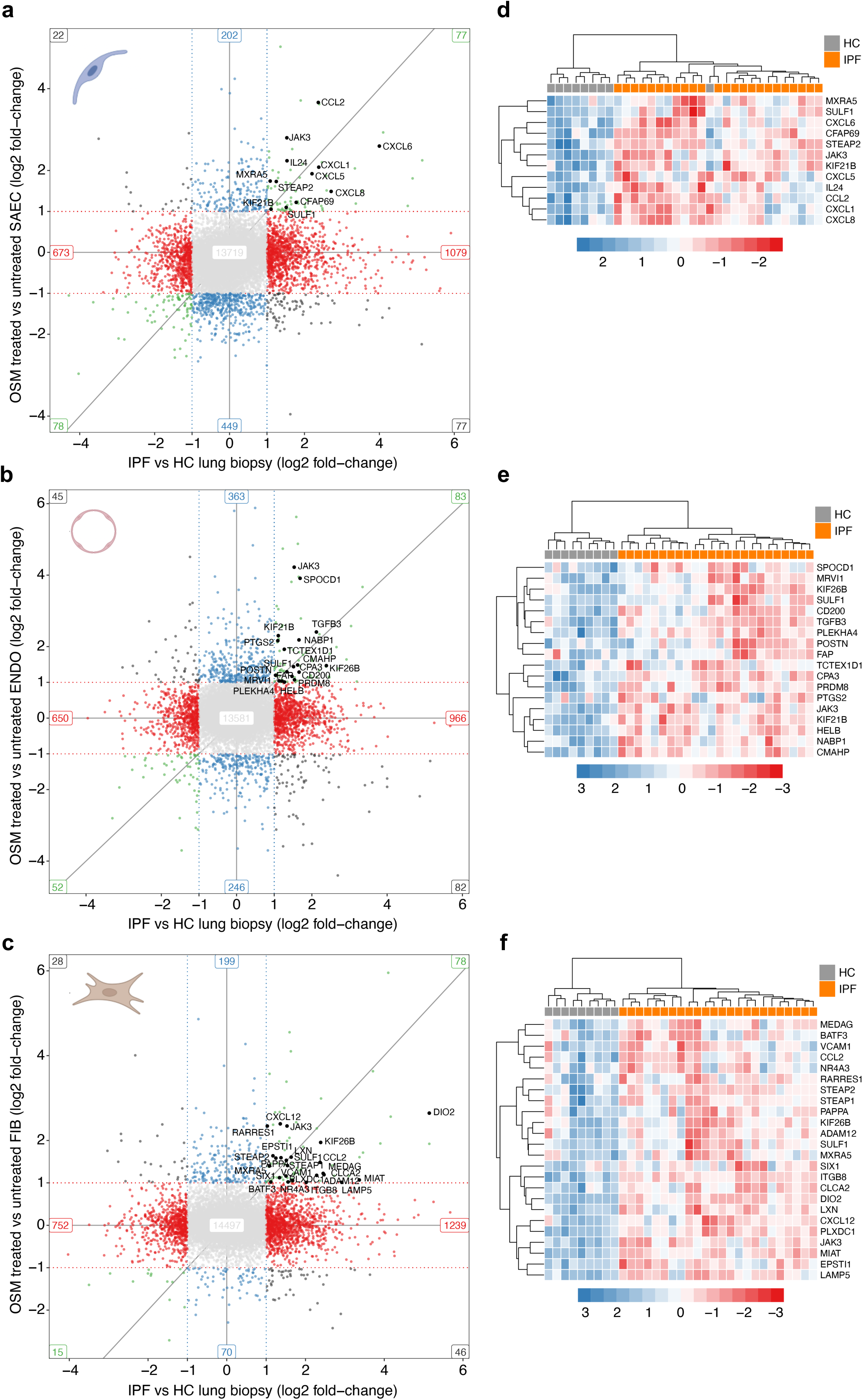
Correlation between OSM-induced responses *in vitro* and observed responses in IPF and SSc-ILD lung tissue *ex vivo*. Primary human SAEC (a), ENDO (b) or FIB (c) were cultured, as described in methods, and stimulated with rhOSM (10ng/ml) for 24hrs. These transcriptional responses are plotted against observed transcriptional responses in IPF lung tissue (left). The common genes identified between *in vitro* stimulated cells and *ex vivo* observed responses in IPF are then plotted in heatmaps for genes significantly upregulated in IPF (FDR P<0.05) to identify concordance between OSM-driven cell type specific response and upregulation in IPF lung tissue (d-f).

**Extended data 10.**
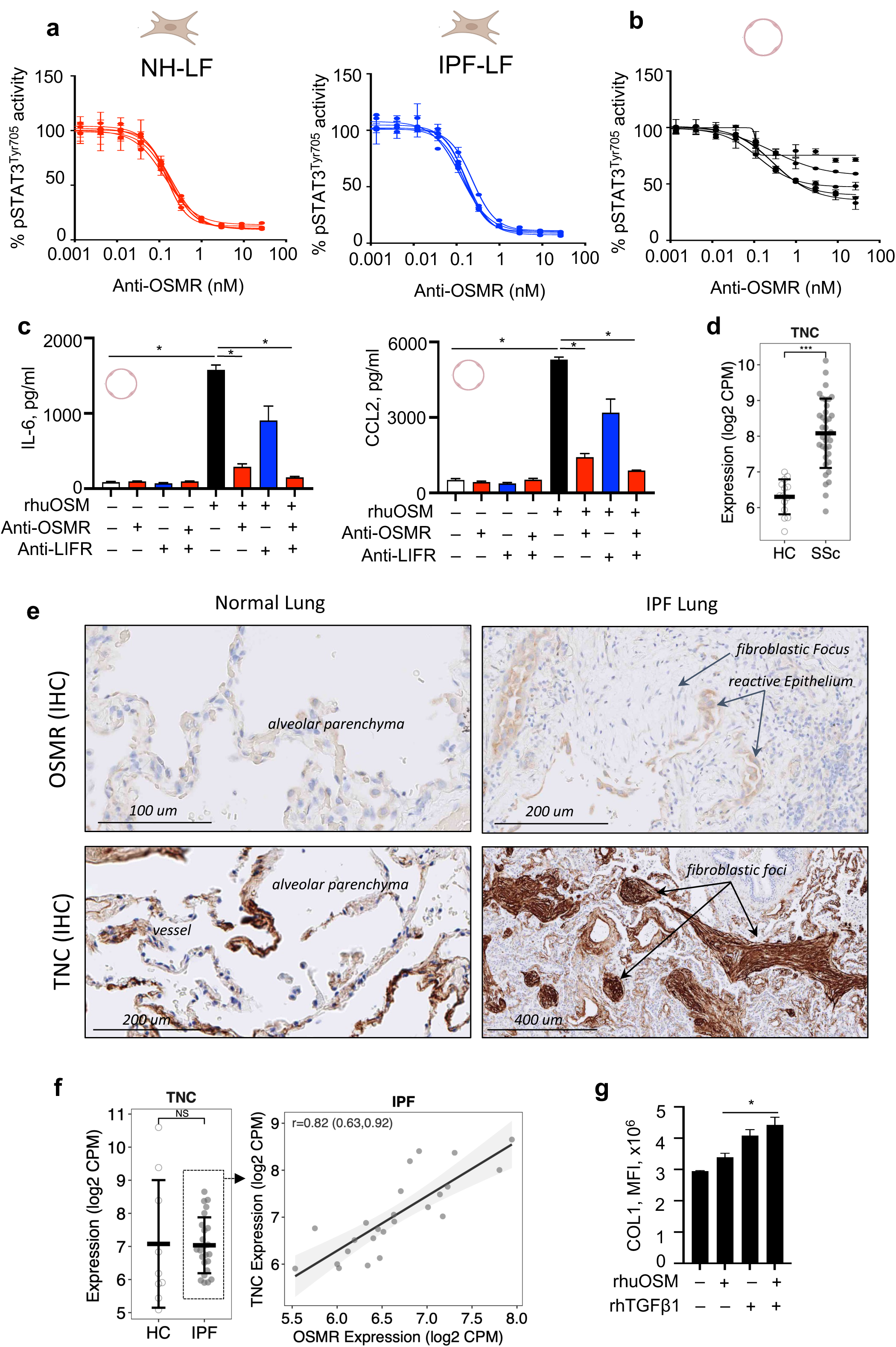
OSMR-dependent activity of OSM in disease-relevant cells. a. Normal primary human lung FIB (NH-LF) or IPF-derived human lung FIB (IPF-LF) were cultured, as described in methods, and stimulated with rhOSM (10ng/ml). Some cells were treated with anti-OSMR, at the indicated concentrations, from 120 mins prior to OSM treatment. Cell lysates were recovered and pSTAT3^Tyr705^ was measured by MSD. Data is expressed as % of remaining pSTAT3^Tyr705^. b. Primary human ENDO cells were cultured, as described in methods and stimulated with rhOSM (10ng/ml). Some cells were treated with anti-OSMR, at the indicated concentrations, from 120 mins prior to OSM treatment. Cell lysates were recovered and pSTAT3^Tyr705^ was measured by MSD. Data is expressed as % of remaining pSTAT3^Tyr705^. c. Primary human ENDO cells were cultured, as described in methods, and stimulated with rhOSM (10ng/ml) for 24hrs with anti-OSMR (50ug/ml) or anti-LIFR (50ug/ml) from 120 mins prior to OSM treatment and throughout. IL-6 and CCL2/MCP1 were measured by luminex in supernatants. P value calculated by t-test. Graphs show mean ± SD. * P<0.05. d. Skin biopsies obtained from healthy controls (n=14) and SSc patients (n=36), participating in the focuSSced clinical trial and *TNC* transcripts measured. Graphs show mean ± SD. *** FDR P<0.0005 e. Normal human lung and IPF lung tissue was stained with anti-human OSMR or anti-human TNC. OSMR+ staining and TNC+ staining is shown. f. Lung biopsies were obtained from healthy controls (n=9) and IPF patients (n=24), RNA-seq was carried out and *TNC* transcripts analyzed (left). TNC levels were plotted against *OSMR* expression (right); linear fit, Pearson correlation, and 95% confidence interval is shown. Graphs show log_2_ normalized CPM, mean ± SD. NS not significant g. Primary human FIB cells were cultured, as described in methods, and stimulated with rhOSM (10ng/ml) or rhTGFβ (1ng/ml), or both rhOSM and rhTGFβ for 72 hrs. Collagen 1 (COL) secretion was stained and assessed using Cell Insight CX7, in the scar-in-a-jar assay, as described in methods. P value calculated by t-test. Graphs show mean ± SD. * P<0.05.

